# Transcriptional deconvolution reveals consistent functional subtypes of pancreatic cancer epithelium and stroma

**DOI:** 10.1101/288779

**Authors:** Jing He, H. Carlo Maurer, Sam R. Holmstrom, Tao Su, Aqeel Ahmed, Hanina Hibshoosh, John A. Chabot, Paul E. Oberstein, Antonia R. Sepulveda, Jeanine M. Genkinger, Jiapeng Zhang, Alina C. Iuga, Mukesh Bansal, Andrea Califano, Kenneth P. Olive

**Affiliations:** Department of Biomedical informatics; Department of Systems Biology; Department of Medicine, Division of Digestive and Liver Diseases; Department of Pathology and Cell Biology; Department of Surgery, Division of GI/Endocrine Surgery; Department of Medicine, Division of Hematology and Oncology; Department of Epidemiology, Mailman School of Public Health; Herbert Irving Comprehensive Cancer Center, Columbia University Medical Center, New York, NY 10032; Department of Computer Science and Engineering, University of California, San Diego; Department of Neuroscience, Icahn School of Medicine at Mount Sinai, New York; PsychoGenics Inc., Tarrytown, New York, USA.

**Keywords:** Pancreatic Cancer, Pancreatic Ductal Adenocarcinoma, Gene Expression, Deconvolution, Laser Capture Microdissection, Subtypes, Stroma

## Abstract

Bulk tumor tissues comprise intermixed populations of neoplastic cells and multiple lineages of stromal cells. We used laser capture microdissection and RNA sequencing to disentangle the transcriptional programs active in the malignant epithelium and stroma of pancreatic ductal adenocarcinoma (PDA). This led to the development of a new algorithm (ADVOCATE) that accurately predicts the compartment fractions of bulk tumor samples and can computationally purify bulk gene expression data from PDA. We also present novel stromal subtypes, derived from 110 microdissected PDA stroma samples, that were enriched in extracellular matrix– and immune–associated processes. Finally, we applied ADVOCATE to systematically evaluate cross–compartment subtypes spanning four patient cohorts, revealing consistent functional classes and survival associations despite substantial compositional differences.

## Introduction

All carcinomas harbor both transformed malignant cells and non-transformed stromal cells, in varying proportions (Aran et al., 2015). Pancreatic ductal adenocarcinoma (PDA) is among the most stroma–rich cancers, with a complex inflammatory microenvironment that typically dominates the tumor parenchyma. Expected to be responsible for over 43,000 deaths per year in the United States, it is a common, aggressive malignancy that responds poorly to therapeutic intervention (Oberstein and Olive, 2013; Siegel et al., 2017). Within the stromal compartment of PDA, diverse fibroblast, myeloid, lymphoid, endothelial and other cell lineages contribute to both pro– and anti–tumor processes, including angiogenesis and epithelial differentiation (Rhim et al., 2014), tissue stiffness (Jacobetz et al., 2012; Provenzano et al., 2012), drug delivery (Olive et al., 2009), and local immunosuppression (Vonderheide and Bayne, 2013). These functions are orchestrated through a host of paracrine signals that pass between and within the epithelial and stromal compartments–communication that is quickly altered upon tissue disruption. Thus, efforts to parse transcriptional programs of PDA should take into account the processes active in both compartments, ideally in an *in situ* context.

Despite extensive genomic characterization (Bailey et al., 2016; Biankin et al., 2012; Jones et al., 2008; Waddell et al., 2015; Witkiewicz et al., 2015), individual DNA mutations have thus far failed to provide confirmed prognostic or theranostic information for PDA. Indeed, only a small fraction of pancreatic tumors is predicted to harbor “druggable” genetic alterations (Bailey et al., 2016; Witkiewicz et al., 2015). As an alternative to genetic biomarkers, transcriptional classifiers for PDA have been explored using bulk tumor samples (Bailey et al., 2016; Collisson et al., 2011; Moffitt et al., 2015). While each study differs in the number of subtypes described, a shared message is that ductal pancreatic tumors include at least two groups distinguished by markers of epithelial differentiation state, with the more poorly–differentiated subtype (i.e. “Basal-like”, “Squamous”, or “Quasi-Mesenchymal”) exhibiting reduced overall survival relative to well-differentiated subtypes (i.e. “Classical” or “Progenitor”). However, the contributions of stromal cells are handled differently in each instance, leading to some debate as to the merits of different proposed subtypes. To clarify this issue, we endeavored to directly profile gene expression from purified neoplastic epithelium and associated stroma isolated from frozen human PDA samples.

Several techniques may be employed to isolate cellular subsets from bulk tissue including magnetic separation, fluorescence assisted cell sorting (FACS), and laser capture microdissection (LCM). The first two techniques rely on population–specific antibodies to separate a suspension of cells following disruption of the tumor. Unfortunately, the extremely fibrotic extracellular matrix of PDA necessitates prolonged enzymatic digestion to achieve a single-cell suspension, during which time transcriptional profiles will be altered. Moreover, PDA diffusely infiltrates the surrounding pancreatic parenchyma (Hruban, 2007) so that even tumor samples enriched by FACS for epithelial markers can include contributions from normal, atrophic, pre-neoplastic, or metaplastic epithelial cells. Laser capture microdissection (LCM) provides a powerful solution, allowing the isolation of relatively pure compartment–specific tissue samples based on morphological features, without disrupting the delicate interplay of intercellular communication. However, performing LCM while maintaining RNA quality suitable for sequencing is technically challenging, costly, and labor intensive. As a result, there are currently few examples of large collections of cancer gene expression profiles derived from LCM samples.

We present here expression profiles of laser capture microdissected malignant epithelium and matched reactive stroma for 66 human pancreatic ductal adenocarcinomas. These data informed the development of a novel computational algorithm for the **A**daptive **D**econ**V**olution **O**f **CA**ncer **T**issue **E**xpression (ADVOCATE), which we used to extend our analyses to external datasets. ADVOCATE may be used to perform two functions on bulk tumor expression profiles. First, it can infer the fractional contribution made by each compartment to the original bulk sample. Second, it can transform a bulk tissue transcriptome into separate “virtual” profiles for each sub–compartment. Though prior approaches exist for related applications (Abu-Alainin et al., 2016; Kuhn et al., 2011; Zhong et al., 2013), they make several assumptions, such as the linear mixing of compartments and the specificity of marker genes, that lead to reduced performance relative to ADVOCATE. Furthermore, a crucial new feature is that ADVOCATE can full infer complete gene expression profiles for modeled subpopulations on a sample–by–sample basis, which may then be input into downstream analytical pipelines.

Given the many cancer-associated processes that are mediated by stromal elements, and the prominence of the tumor stroma specifically in PDA, molecular subtyping of stromal signaling programs may offer a distinct and useful counterpoint to epithelial classification. A prior study utilized an elegant computational approach to indirectly infer stromal subtypes based on a limited set of stroma–associated expression factors (Moffitt et al., 2015). Here we provide a novel stromal classification derived from a total of 110 experimentally purified stromal profiles. Two prominent subtypes were apparent, enriched for extracellular matrix associated pathways (ECM–rich) and immune/cytokine associated pathways (Immune–rich), respectively. Critically, application of this novel, stroma–specific gene signature to virtual stroma profiles generated by ADVOCATE from bulk tumor samples led to the consistent identification of the same functional subtypes across three large external PDA gene expression datasets; similar consistency was also observed using an existing epithelium–specific classification system. Notably, a meta-analysis of all four cohorts revealed a partial association of the ECM–rich stroma subtype with the Basal–like epithelial subtype. Finally, we demonstrated that incorporating molecular subtypes from both the epithelial and stromal compartments into a combined classification led to the identification of subtypes that are strongly associated with patient survival across multiple cohorts.

## Results

### Transcriptional profiling of pancreatic cancer epithelium and stroma

To study the separate transcriptional programs of intact pancreatic tumor epithelium and stroma, we optimized a robust protocol for maintaining RNA integrity during laser capture microdissection of frozen tumor tissues, yielding total RNA suitable for library preparation and RNA sequencing. We first applied this LCM–RNA–Seq technique to 60 primary PDA specimens that were harvested and frozen intraoperatively by the Columbia University Tumor Bank in collaboration with the Columbia Pancreas Center (see Tables S1,2 for patient characteristics). For each tumor, we generated paired gene expression profiles from the malignant epithelium and nearby reactive stroma, as distinguished by cell morphology (Figure 1A). Extensive quality control metrics confirmed the high quality of resulting RNA libraries (Figures 1B,C and Figures S1A-D) (Adiconis et al., 2013; Shanker et al., 2015). Critically, samples from the two compartments separated spontaneously along the first component of a Principal Component Analysis (PCA) with virtually no overlap (Figure 1D), and were distinguished by expression of established marker genes for epithelial cells (KRT19, EPCAM, CDH1) versus markers of various stromal cell types, including leukocytes (PTPRC, CD4, CD163), endothelial cells (VWF, ENG, CDH5), and cancer associated fibroblasts (CAFs) (ACTA2, DCN, FAP) (Figure 1E). We observed that technical variance was substantially lower than biological variance (Figures S1E-F) and found that different malignant areas captured from a single tumor clustered closely, suggesting that the intra-tumoral transcriptional heterogeneity of that tumor was less than the inter-tumoral heterogeneity of PDA (Figures S1G-H). In summary, our analysis shows that LCM–RNA–Seq produces robust, genome–wide, compartment–specific gene expression profiles.

**Figure 1.**
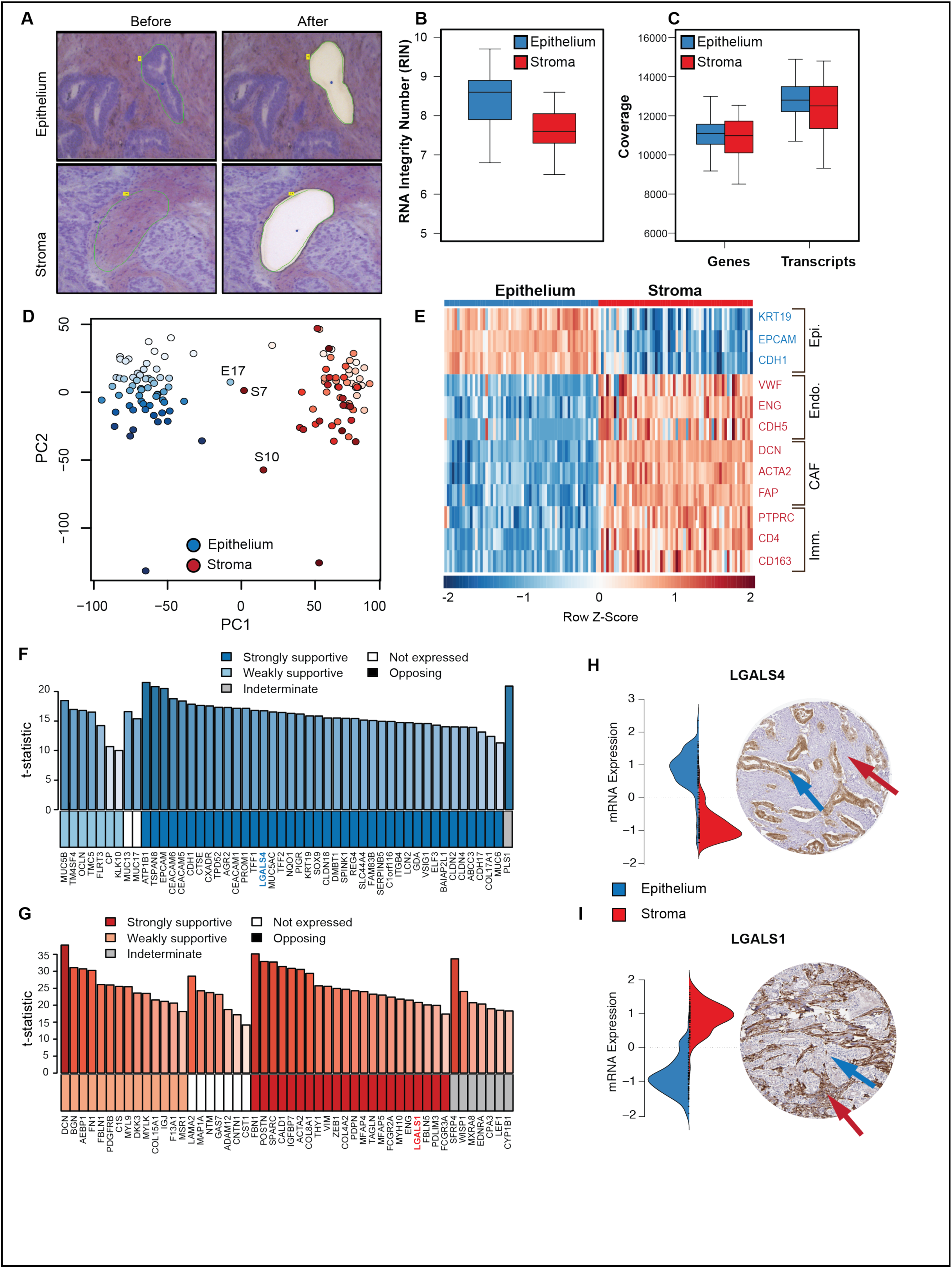
Compartment–specific gene expression profiling of pancreatic tumors. (**A**) Images of Cresyl Violet stained human PDA frozen sections before and after laser capture microdissection of malignant epithelial and adjacent stromal cells. (**B**) RIN values for RNA samples derived from the indicated compartment (N = 60 each). (**C**) Number of genes and transcripts detected at > 1 FPKM in the samples from (B). (**D**) Principal component analysis of the profiles from (C). 60 pairs of epithelial and stromal LCM samples. Color graduation shows pairing of samples from the same tumor. Three samples discussed later are labeled. (**E**) Heatmap showing the differential expression of marker genes for the indicated cell types in each sample. (**F,G**) Validation of genes predicted as epithelium–specific (F) or stroma–specific (G) based on mRNA expression. Bar height and color shading reflects the certainty (t-statistic) of differential expression. The box color below each bar summarizes results of immunohistochemistry on PDA sections from the Human Protein Atlas (HPA). IHC staining pattern was categorized as strongly or weakly supportive of the predicted compartment (blue/red), indeterminate (grey), absent (white), or opposite the predicted pattern (black). (**H**) An example epithelium–specific gene, LGALS4, showed a protein staining pattern that was strongly consistent with its mRNA expression (at left). Blue and red arrows indicate PDA epithelium and stroma, respectively. (**I**) An example stroma–specific gene, LGALS1, was expressed exclusively in the tumor stroma.

We next examined the most differentially expressed genes between the epithelium and the stroma (Table S3) and compared their expression to immunohistochemistry for the corresponding proteins in The Human Protein Atlas (HPA) pathology database (Pontén et al., 2008). We restricted our analysis to proteins for which the highest-quality antibodies were available (n= 321), based on established HPA criteria (Table S4). Of these, we evaluated the immunostaining patterns for the 50 genes whose LCM–RNA–Seq expression was most differentially expressed for each compartment, examining a minimum of six PDA samples per tested protein. This analysis yielded confirmatory staining patterns for 47 of 50 epithelial proteins and 36 of the 50 stromal proteins (Figures 1 F, G, Table S5). For example, Figures 1 H, I show two members of the galectin protein family, LGALS4 and LGALS1, with inverse staining patterns in the two compartments, consistent with our predictions. Critically, none of the proteins were found expressed in a pattern opposite that predicted; most genes lacking supportive staining were simply not detected, perhaps due to post–translational regulation. Thus, through the use of LCM–RNA–Seq, we compiled a rich resource of compartment–specific genes that may be of use as novel markers for the pancreatic cancer field.

### A framework to deconvolve compartment–specific gene expression profiles from bulk data

Multiple large–scale gene expression datasets for PDA have been reported and each has provided important contributions to our understanding of the disease. However, it has been challenging to make comparisons between these datasets due to differences in expression profiling platforms, inclusion criteria, sample preparation, and other technical details. The availability of experimentally–purified, compartment–specific, paired expression profiles offered a unique opportunity to unify these resources by using transcriptional deconvolution to remove the noise introduced by variable tissue composition in each dataset. In contrast to many prior optimization approaches (i.e. quadratic programming), we utilized a machine-learning approach to train a probabilistic algorithm (ADVOCATE), consisting of four steps, as outlined in Figure 2A: 1) ADVOCATE fits a Gaussian-mixture model on the collection of normalized pairs of expression profiles to infer the expression distribution of each gene in each compartment. 2) When subsequently provided with a bulk gene expression profile, the expression of each gene in the bulk sample is compared with the expression distribution inferred in Step 1, to derive a compartment fraction prediction for each detected gene. 3) These fractions are integrated across all genes to infer a global compartment fraction for the bulk sample. 4) Finally, ADVOCATE combines the information from steps 1, 2, and 3 to infer compartment–specific virtual profiles, theoretically recapitulating the results of more–costly and laborious LCM–RNA–Seq. While we focus here on the epithelial and stromal compartments of pancreatic cancer, there is no technical limitation for the number of compartments that can be modeled provided that distinct, matched, compartment–specific expression profiles are available.

**Figure 2.**
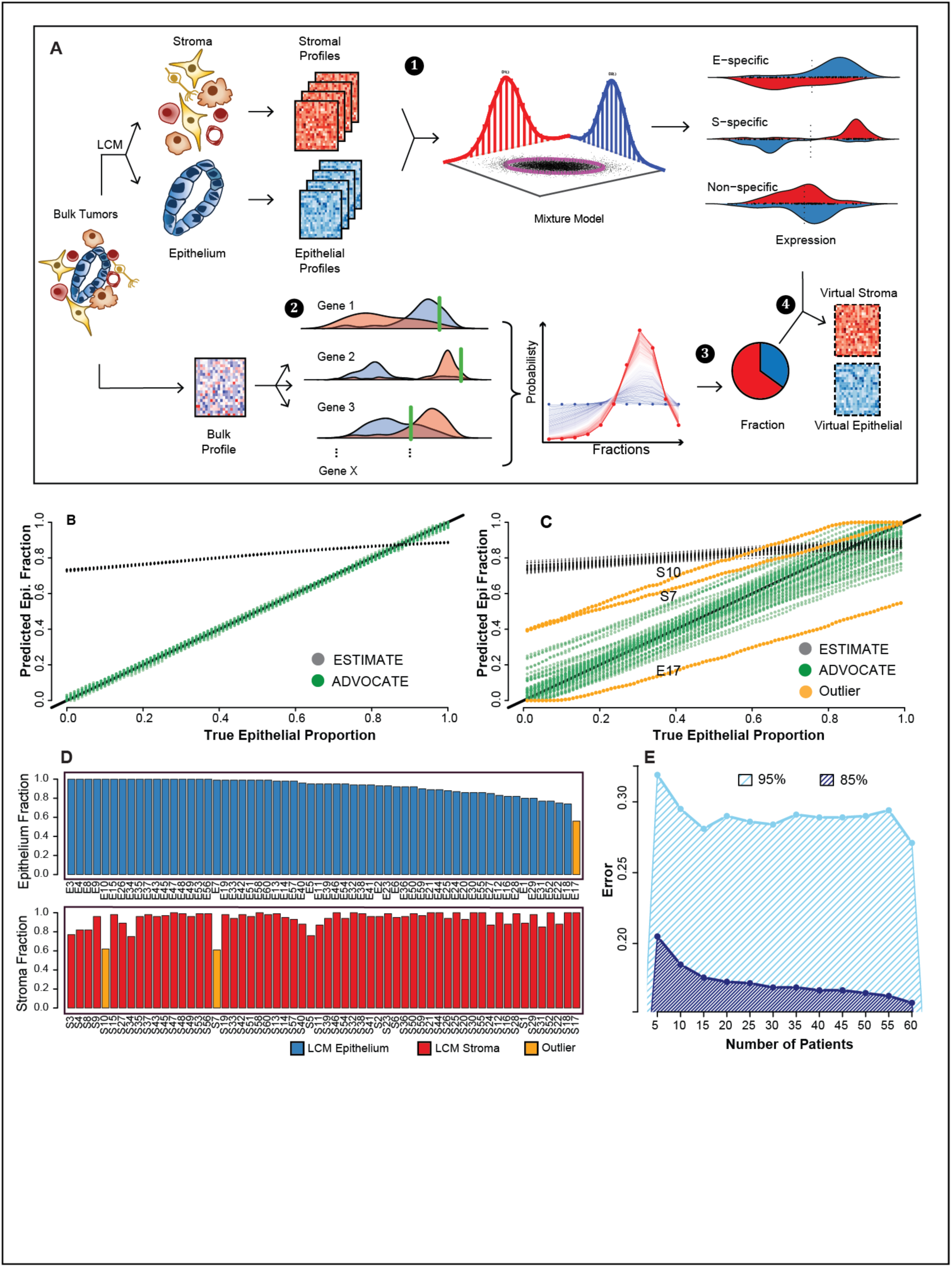
ADVOCATE framework and computational validation. (**A**) Outline of the ADVOCATE framework where LCM–RNA–Seq profiles are used to model genome–wide gene expression distributions for each gene across compartments. Bulk tissue profiles can be deconvolved using compartment–specific, gene–wise probabilities, yielding two products: i) predicted compartment fractions in the bulk tumor and ii) virtual gene expression profiles for the epithelium and stroma of that sample. Numbers indicate steps described in main text. (**B**). Compartment fraction analysis performed with ADVOCATE (green) or ESTIMATE (black) on a computationally generated (synthetic) set of epithelial and stromal profiles mixed together *in silico* in varying proportions. (**C**) Compartment fraction analysis performed with ADVOCATE (green/yellow) or ESTIMATE (black) on a semi-synthetic set of samples generated by mixing actual experimental pairs of LCM expression profiles together in varying proportions, for each of 60 tumors. The yellow lines highlight three outlier samples. (**D**) Leave-one-out cross validation of compartment fraction using ADVOCATE where the epithelial (blue) and stromal (red) fractions, are predicted for each individual pair of LCM–RNA–Seq profiles after training the model using the remaining 59 pairs. The outliers from (C) are indicated as yellow bars. (**E**) Power analysis for training the ADVOCATE algorithm. Plot shows 95% (dark blue) and 85% (light blue) percentile of prediction errors relative to sample sizes based on LOOCV analysis. Errors decreases quickly as sample size increases, stabilize after ∼ 20 samples, and reach a minimum at 60 samples.

### Computational validation of ADVOCATE performance

We carried out a series of *in silico* and experimental assays to validate ADVOCATE’s performance. First we generated artificial pure epithelial and stromal gene expression profiles by randomly sampling from the density of expression for each gene across the LCM–RNA–Seq profiles for each compartment (see Methods). These were then mixed together *in silico* in varying proportions to create synthetic bulk samples with different compartment fractions. When ADVOCATE was trained on these synthetic compartment–specific samples, it predicted the compartment fractions of the synthetic bulk samples with error rates of less than 3% (Figure 2B). By comparison, the ESTIMATE algorithm (Yoshihara et al., 2013), which is commonly used for compartment fraction analysis of many cancer types, significantly and systematically over–estimated the epithelial compartment fraction, independent of the actual compartment mixture (black dotted line in Figure 2B).

Next we generated a more realistic set of “semi–synthetic” bulk samples by computationally mixing profiles from actual LCM–derived epithelial and stromal sample pairs, in varying proportions (Figure 2C). Again, ADVOCATE predicted the compartment mixture rate with very high accuracy (>90%) for the majority (∼80%) of the samples. In a complementary analysis, we estimated compartment–specific LCM sample purity by leave–one–out cross-validation (LOOCV), whereby the composition of each individual LCM sample was estimated after training the model on the remaining 59 pairs. Figure 2D shows that ADVOCATE correctly predicted a very high (>90%) epithelial and stromal composition for the majority of compartment–specific LCM–derived samples (73%), as expected, with the residual error likely due to biological and technical variability in the LCM–derived sample profiles.

Interestingly, the same three outlier samples with >20% error were detected in the analyses from both Figure 2C and 2D (orange lines/bars). Careful histopathological examination of these samples revealed that one (E17) was poorly differentiated and therefore may plausibly exhibit a more stroma–like signature (Figures S2A-B). A second (S10) had large areas of highly cellular stroma intermixed with fibrotic regions, which could plausibly lead to a more epithelial–like stroma signature (Figure S2C-E). No obvious pathological distinctions were apparent in the third sample (Figure S2F) suggesting either imprecise microdissection or an unusually high level of epithelial delamination into the stroma (Rhim et al., 2012). These findings were consistent with the clustering of these three samples near the interface between epithelial and stromal samples by PCA (Figure 1D). Together, the results provide evidence of the robustness of ADVOCATE in predicting bulk tumor composition from synthetic data and its resilience when applied to more realistic experimental data. A power analysis performed using the LOOCV technique found that experimental error diminished substantially as the size of the training set exceeded 20 tumors, indicating fairly modest sample requirements for training an implementation of ADVOCATE (Figure 2E).

### Experimental validation of compartment fraction prediction with ADVOCATE

We next assessed ADVOCATE’s ability to predict the composition of samples whose compartment fraction had been experimentally assessed and compared it to four other published algorithms: ESTIMATE, deconRNAseq (Zhong et al., 2013), PSEA (Kuhn et al., 2011), and DSA (Zhong et al., 2013). We began by analyzing gene expression profiles from pure epithelial and stromal tissues isolated by LCM on PDA samples (Figures 3A, S3A-B). Among the five algorithms, ADVOCATE predicted the highest epithelial fraction in microdissected epithelial samples, and the second-lowest epithelial fraction in stromal samples; the deconRNAseq algorithm predicted lower epithelial fractions in both the stromal and epithelial datasets. We then used ADVOCATE to predict the compartment fractions of the CUMC bulk samples for which nuclei counting was performed (Figure S3C) as well as bulk PDA samples form TCGA, which are annotated with DNA-based purity estimates using the ABSOLUTE algorithm (Figure S3D). Among the five algorithms, ADVOCATE predicted the highest purity for the respective compartment in the LCMsamples, and scored closest to the respective reference standards for the CUMC and TCGA bulk PDA samples (nuclei count and ABSOLUTE prediction, respectively) (Figure S3E).

**Figure 3.**
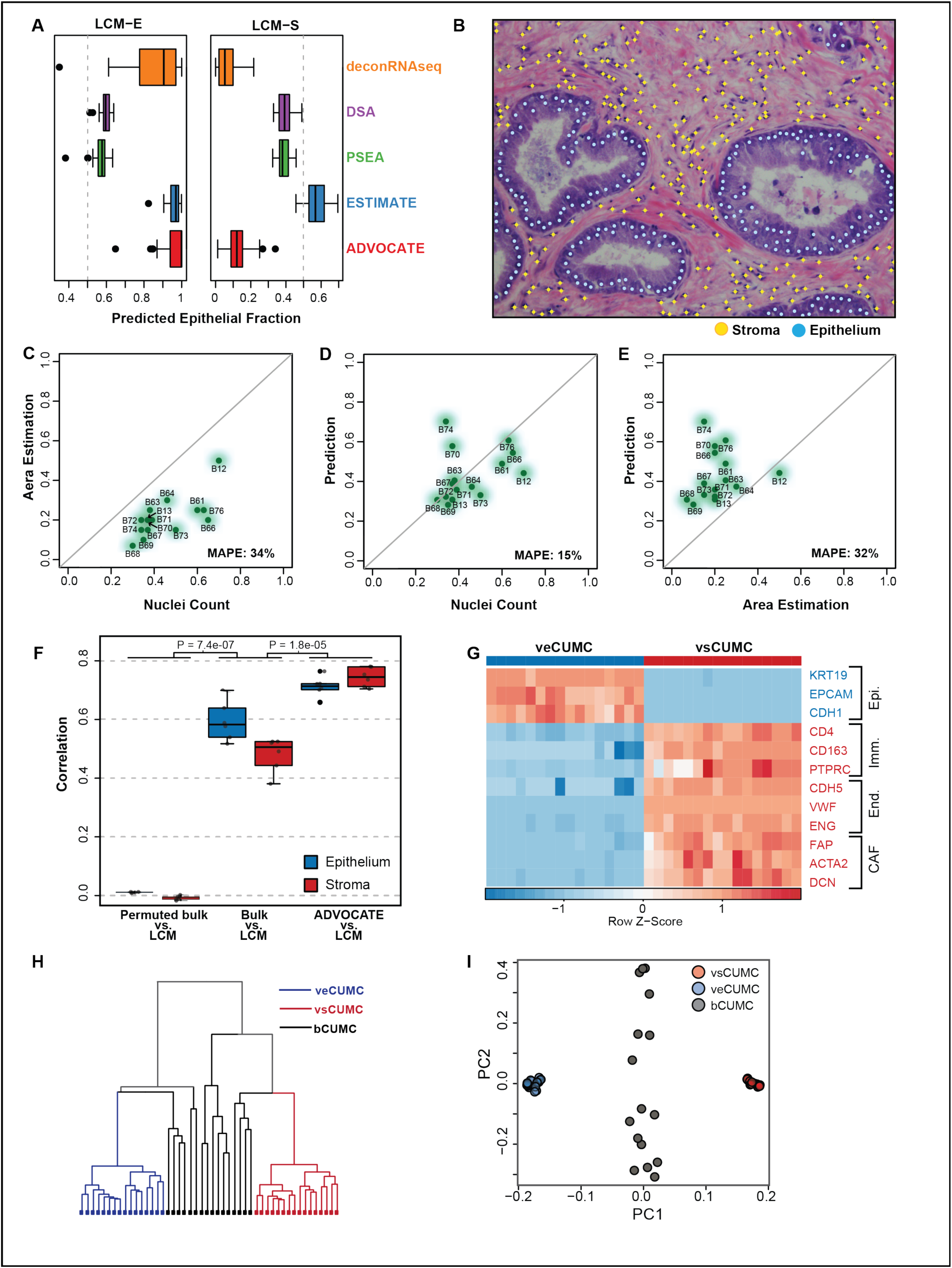
Experimental validation of ADVOCATE. (**A**) Predicted compartment fractions for CUMC epithelial (left) and stromal (right) LCM–RNA–Seq profiles using ADVOCATE (red), ESTIMATE (blue), PSEA (green), DSA (purple) and deconRNAseq (orange). (**B**) Example of nuclei counting for the epithelial (teal dots) and stromal (orange dots) compartments. Nuclei per compartment were divided by the total to yield the compartmental fraction. (**C-E**) Comparison between tumor epithelium content estimated by (**C**) nuclei counting vs. area estimation, (**D**) area estimation vs. ADVOCATE, and (**E**) nuclei counting vs. ADVOCATE for 15 tumors from the Columbia University Medical Center (CUMC) collection. MAPE denotes Mean Absolute Percentage Error as a measure of prediction accuracy. (**F**) Correlation of normalized gene expression between experimental LCM–RNA–Seq profiles and: permuted bulk profiles (left, negative control); bulk expression profiles (middle); and virtual epithelial and stroma profiles purified from bulk using ADVOCATE (right). In each pair, blue indicates epithelium and red indicates stroma. (**G**) Heatmap showing expression of indicated marker genes in virtual epithelial and stromal profiles derived from the 15 bulk tumors described in (C). (**H**) Hierarchical clustering and (**I**) PCA of 15 bulk CUMC tumors and their virtual compartment–specific derivatives.

Carcinoma–derived cell lines are often used to provide a pure epithelial reference for gene expression studies. We next used ADVOCATE to predict the compartment fractions of 40 PDA cell lines in the Cancer Cell Line Encyclopedia (CCLE)(Barretina et al., 2012). As expected, all PDA cell lines were scored as predominantly epithelial with a mean epithelial fraction of 94% (Figure S3F). However, nine lines were predicted to have >10% stromal fraction. This should not be interpreted to imply the presence of stromal cells in these lines. Rather, this result likely indicates either that the original tumor was poorly differentiated or that the cells underwent epithelial–to–mesenchymal transition (EMT) during adaptation to tissue culture conditions, since stromal signatures are enriched in mesenchymal genes (Ross et al., 2000). Consistent with that interpretation, sarcoma and mesothelioma cell lines (n = 28) were predicted to be significantly more “stromal” than PDA cell lines (P = 2.30×10^−25^, Kolmogorov-Smirnov (KS) test, Figure S3G). This finding highlights potential issues in the common practice of using epithelial tumor cell lines indiscriminately as a “pure” epithelial reference for purity assessments.

Finally, to directly test the ability of ADVOCATE to predict compartment fractions from bulk tissue, we performed RNA–Seq on bulk tissue from 15 PDA samples and compared compartment fraction predictions to histopathological assessments. Bulk tumor samples may also contain varying amounts of other tissue types (such as normal or atrophic pancreas, lymphoid aggregates, nerve plexus, and blood vessels) that are routinely found intermixed with frank carcinoma in pancreatic tumors. To account for this, we incorporated a third “Residuals” compartment into this analysis, comprising genes with a low expression probability in both epithelial and stromal compartments (see Methods). We note that multiple studies have reported discrepancies between tumor purity estimates from pathology review and those from molecular analyses (Carter et al., 2012; Song et al., 2012; Yoshihara et al., 2013), either as a result of technical limitations or of sampling different areas of the tumor for the respective analyses. To minimize the latter effect, we assessed the histology of tissue sections immediately adjacent to those used for LCM-RNA-Seq. Two blinded, independent histopathology assessments of tumor composition were performed on hematoxylin and eosin (H&E) stained frozen tissue sections. First, the areas of epithelium, stroma, and other tissue were estimated by a gastrointestinal pathologist. Second, individual nuclei were counted in the epithelial and stromal compartments for multiple representative tumor areas (Figure 3B). We found that tumor content was highly correlated between the two measures (ρ = 0.77), with the area assessment showing lower epithelial content than nuclei counting (Figure 3C). ADVOCATE’s predictions tracked very well with nuclei count assessment, with a mean absolute percent error (MAPE) of 15% (Figure 3D). However, it systematically overestimated tumor content, relative to area assessments (MAPE = 32%, Figure 3E), perhaps suggesting that nuclei count better reflect gene expression contributions from distinct compartments. Overall, ADVOCATE yielded the lowest MAPE of the five algorithms assessed (Figure S3E).

### Experimental validation of virtual expression profiles generated from bulk tumors

The second function offered by ADVOCATE is to extract compartment–specific gene expression data from bulk samples (referred to as “virtual profiles” hereafter). To assess the accuracy of virtual profiles, we performed both bulk RNA-Seq and LCM–RNA–Seq on six tumor samples. We then determined the correlation between the LCM profiles and their bulk counterparts, before and after deconvolution (Figure 3F). Without deconvolution, there was only modest correlation with ρ = 0.59 and ρ = 0.48 for epithelium and stromal LCM comparisons, respectively. By contrast, following deconvolution with ADVOCATE, the correlation between experimental LCM and virtual profiles increased significantly to ρ = 0.71 for the epithelium and ρ = 0.74 for the stroma (P = 1.8×10^−5^ combined, two-sample KS test). Critically, expression of lineage–specific markers in virtual epithelial and stromal samples closely tracked with the results of experimental microdissection (Figure 3G vs. Figure 1E). Indeed, both hierarchical clustering and PCA showed divergence of virtual epithelial and stromal profiles relative to the intermediate clustering of bulk profiles (Figures 3H, I). Taken together, these data demonstrate that ADVOCATE is effective in deconvolving compartment–specific expression profiles from bulk tissue.

Finally, we sought to assess the general suitability of ADVOCATE for deconvolving cancer types by making use of a published array expression dataset derived from laser capture microdissected breast cancer samples (Oh et al., 2015) (Figures S4A, B). We trained an implementation of ADVOCATE on a subset of these samples and then performed LOOCV analysis, finding the predicted fraction to be >80% in the majority of samples for both compartments (Figure S4C). We then applied this model to the remaining samples from the dataset, including samples that clustered with the training samples as well as some expected to have intermediate compartment fractions, based on their position in the PCA (yellow points in Figure S4B). As expected, most of the samples from this test were predicted to have high purity, but compartment fractions were lowest in the intermediate samples (Figure S4D). Finally, we used this implementation of ADVOCATE to predict the compartment fractions of an independent LCM RNA-seq breast cancer dataset (N=36, Agilent platform, GSE68744) and compared these results to purity predictions from ESTIMATE (Figure S4E). Critically, ADVOCATE successfully predicted quite pure fractions for the relevant compartment on these independent LCM samples. By contrast, ESTIMATE predicted lower purity in both compartments, particularly in epithelium samples. These results demonstrate both the generalizability of the ADVOCATE algorithm across tumor types when provided appropriate training samples, and the suitability of ADVOCATE for use with different gene expression platforms.

### Compartment fraction analysis reveals distinct compositions of public PDA datasets

Three prior studies have presented expression–based classification systems, derived from large collections of bulk PDA profiles, that relate to biological properties or outcome (Bailey et al., 2016; Collisson et al., 2011; Moffitt et al., 2015). In each study, a subtype exhibiting molecular characteristics consistent with a poorly differentiated state (termed “Quasi-Mesenchymal”, “Basal-like”, or “Squamous”, respectively) was shown to have poor overall survival, relative to a subtype exhibiting a signature reflective of pancreatic tissue origin (“Classical” or “Progenitor”), with varying numbers of additional groups also presented. Moffitt et. al. further parsed molecular subtypes from the tumor stroma using non-negative matrix factorization (NMF, a statistical approach to separating the constituent parts of an object. See Discussion.)(Lee and Seung, 1999). Each of these reports has contributed to our basic understanding of pancreatic cancer biology, but a systematic evaluation of compartment–specific contributions is needed to identify subtypes that are broadly applicable across biologically and technically heterogeneous PDA patient cohorts.

We used the compartment fraction analysis function of ADVOCATE to analyze pancreatic cancer gene expression profiles from three independent cohorts: (a) UNC Chapel Hill (UNC, n = 125), (b) the International Cancer Genome Consortium (ICGC, PACA-AU RNA-Seq dataset, n = 93), and (c) The Cancer Genome Atlas (TCGA, PAAD dataset, n = 137) (see Methods for inclusion criteria). We reasoned that there are three potential sources of relevant heterogeneity in these data: 1) differences in the epithelium-to-stroma ratio in areas of frank carcinoma; 2) variation in the extent of non-tumor tissues (normal pancreas, pancreatitis, lymph nodes, etc.) included in the bulk sample; and 3) technical differences (e.g. expression platform, library preparation method, etc.). We reasoned that the contributions of the latter two sources will be largely captured by the “Residuals” component of the 3–compartment implementation of ADVOCATE, since they would lead to gene distributions distinct from those found in the pure LCM datasets. Thus, subtracting the Residuals component serves as a means to control for technical and biological heterogeneity independent of true compartment fraction. Using this approach, we found that the epithelial and stromal fractions varied significantly between the cohorts with 46%, 67% and 55% epithelium for the ICGC, UNC and TCGA cohorts, respectively (p < 0.001, one-way ANOVA) (Figure 4A). This highlights critical differences in composition between tumor collections curated with different inclusion criteria or enrichment practices.

**Figure 4.**
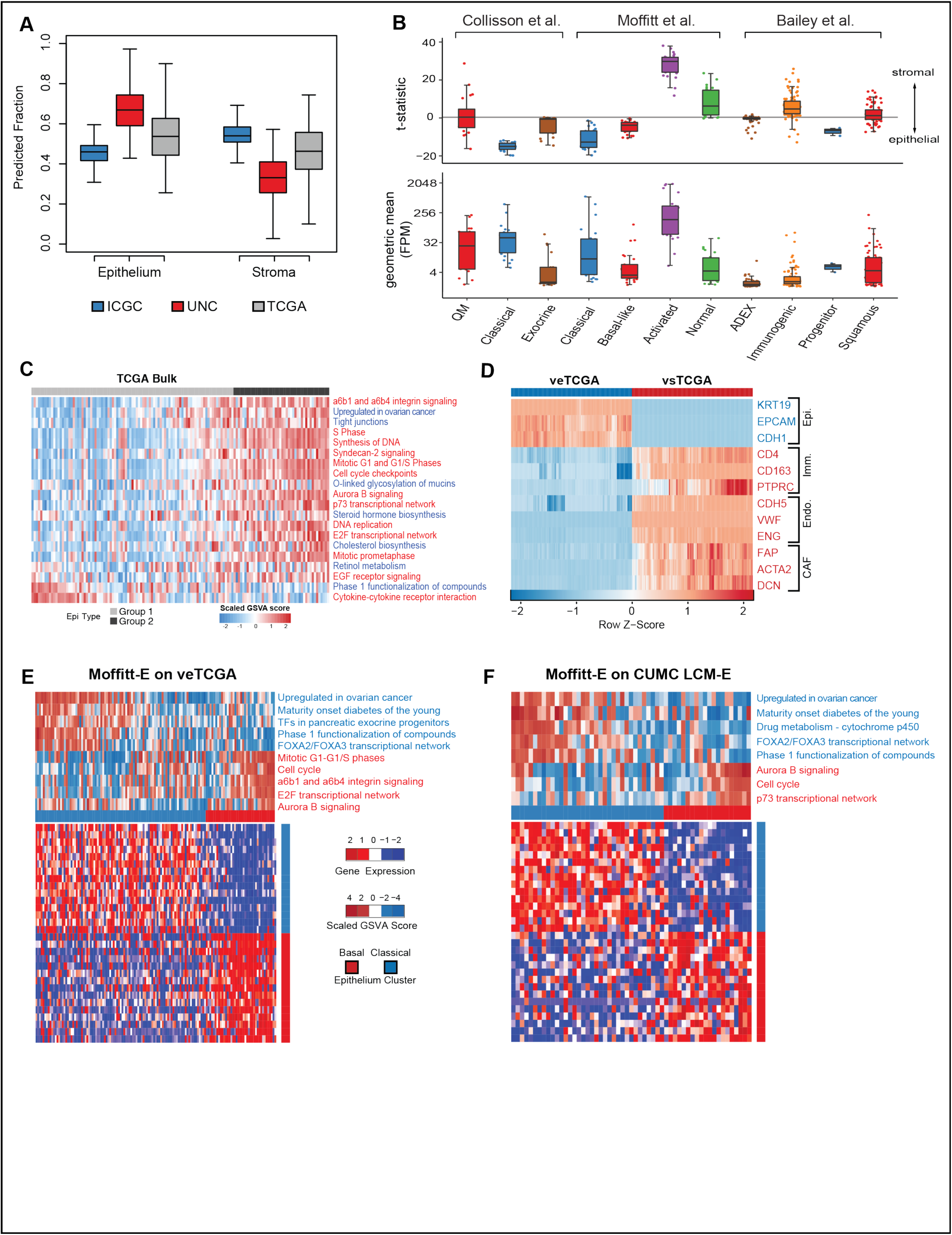
Analysis and classification of pancreatic tumor cohorts and classifiers. (**A**) Compartment fraction analysis of pancreatic tumors from the ICGC (blue), UNC (red), and TCGA (grey) cohorts. The 3-compartment implementation of ADVOCATE was used for fraction estimation to reduce noise from technical and non-tumor sources, and the epithelial and stromal estimate were then scaled to sum up to 1. (**B**) Analysis of genes comprising three published classifiers for PDA using LCM–RNA–Seq data. Upper panel depicts the distribution of t-statistics for each of the indicated classifier gene lists that were calculated by paired differentially expressed gene analysis between the epithelium and stroma. Positive values indicate stromal enrichment. Lower panel depicts the geometric mean expression in fragments per million mapped fragments (FPM) for the genes from each classifier gene list across all LCM–RNA–Seq samples. (**C**) Heatmap depicts the enrichment of each TCGA bulk sample for the indicated gene sets. Gene sets indicated functions associated with Basal-like (red) or Classical (blue). (**D**) Heatmap depicts the enrichment of expression of indicated marker genes in deconvolved virtual epithelial and stromal profiles from the TCGA cohort. (**E**) Hierarchical clustering TCGA bulk tumors (black) and their virtual epithelial (blue) and stromal (red) derivatives. (**F,G**) Heatmaps of the top 30 differentially expressed genes between two groups that were obtained by clustering virtual epithelial TCGA (F) and CUMC (G) profiles using the Moffitt–E classifier. GSVA scores for indicated gene set are represented at top. Tumors with Basal–like (red bar) or Classical (blue bar) traits, respectively, can be detected in both virtual tumor cohorts.

We next focused on the expression level and compartment–specificity of the genes used by each of the three published PDA classification systems (Figure 4B) (Bailey et al., 2016; Collisson et al., 2011; Moffitt et al., 2015). We began by remapping the Columbia University Medical Center (CUMC) dataset using the Ensembl GRCh37 gene annotation in order to enable comparisons to the ICGC dataset, which uses this annotation. This resulted in 22% more genes being called than with the NCBI annotation, and had a particularly strong impact on the many recombined immunoglobulin genes that contribute to the Bailey et. al. Immunogenic subtype (Tables S17-20). Notably, we observed that the genes used to define the Classical, Basal-like, and Progenitor subtypes were heavily weighted towards epithelium–specific expression. Conversely, those used to define the Activated, Normal and Immunogenic subtypes were weighted towards the stroma. The Quasi-Mesenchymal (QM) and Squamous gene sets were well–expressed and represented a mixture of epithelial and stromal identity, consistent with a more poorly differentiated state. Finally, the majority of genes that define the Exocrine and ADEX subtypes exhibited very low expression in the LCM-RNA-Seq datasets, suggesting that their expression in bulk tissue is derived from cell types largely absent from our microdissected samples. Together, these data provide insight into the cellular compartments that contribute to molecular gene signatures built from bulk tumor tissue samples.

### Transcriptional deconvolution improves functional classification across cohorts

An important feature of a robust classification system is its capacity to identify functionally similar groups across independent datasets. A gene signature can stratify groups of samples that are functionally unrelated due to variation between the cohorts. For example, application of the epithelial signature from Moffitt et. al. (hereafter referred to as Moffitt–E) to bulk profiles from TCGA identifies two groups of samples. However, gene set variance analysis of the two TCGA groups finds intermixing of the functions associated with the Basal–like and Classical groups (Figures 4C). With the finding that compartment fraction varies substantially between different cohorts, we wondered whether removing the expression contributions of stromal cells would aid classification efforts.

To address this challenge, we used the deconvolution function of ADVOCATE to generate virtual epithelial and stromal expression profiles from the bulk samples of each PDA cohort (producing datasets vUNC, vTCGA, and vICGC). In each case, the resulting virtual profiles were clearly distinguished by established cell–specific marker genes (Figure 4D and Figure S5A, B). Notably, bulk samples were distributed between the corresponding virtual epithelium and stroma samples by hierarchical clustering (Figures S5 C–E).

To address whether deconvolution improved cross–cohort consistency of molecular classifiers, we processed the <u>v</u>irtual <u>e</u>pithelial profiles from each cohort (veUNC, veTCGA, and veICGC) using the Moffitt–E signature. Notably, deconvolution of TCGA data led to a realignment in functional associations, resulting in two groups that could be clearly recognized as Basal-like and Classical, based on enriched gene sets (Figure 4E). Similar functional associations were observed when the Moffitt–E classifier was applied to our experimentally–derived epithelial LCM–RNA–Seq (CUMC LCM–E) dataset (Figure 4F). We also noted that application of the Moffitt–E classifier to the veICGC dataset revealed excellent alignment with the pancreatic progenitor and squamous subtypes described by (Bailey et al., 2016) (SMC = 0.91) (Table S6). Together, these data indicate that removal of stromal expression data from bulk tumor datasets can improve the functional similarity of the groups identified by classification systems.

### Identification of Immune-rich and ECM-rich subtypes of PDA stroma

The profuse stromal desmoplasia of pancreatic ductal adenocarcinoma is a defining feature of this malignancy. Previously, Moffitt and colleagues used NMF (Lee and Seung, 1999) on a large number of bulk PDA expression profiles to infer a PDA stromal classifier that identified two subtypes, designated “Activated” and “Normal” (Moffitt et al., 2015). These subtypes were interpreted to reflect the biology of cancer associated fibroblasts, based on the inferred contributions of activated myofibroblasts versus quiescent pancreatic stellate cells. While this finding both aligns well with known PDA biology and was shown to be clinically relevant with respect to outcome association, we were interested in capturing expression signals from the stroma as a whole, including the many myeloid, lymphoid, endothelial and other cell types that are commonly present in the PDA microenvironment. To this end, we expanded the stromal LCM–RNA–Seq cohort described above to include samples from a total of 110 unique patients. NMF with consensus clustering identified two prominent molecular subtypes among these samples. Clear functional identities were established for these subtypes using gene set variance analysis (GSVA), leading to their designations as: an “Immune–rich” group characterized by numerous immune and interleukin signals; and an “ECM–rich” group, characterized by numerous extracellular matrix–associated pathways (Figure 5A). We next extracted a gene signature distinguishing these two stromal subtypes, making use of the compartment specificity analysis described above to filter for stroma-specific genes (see Supplementary Methods and Supplementary Tables 22 – 24). Application of this signature to the virtual stroma profiles yielded two prominent groups each for the UNC, ICGC, and TCGA cohorts (Figures 5B–D). Critically, in each cohort, the two groups were again characterized by their enrichment for gene sets associated with ECM deposition or immune processes, indicating a robust and consistent performance of this new, stroma-specific “CUMC–S” signature.

**Figure 5.**
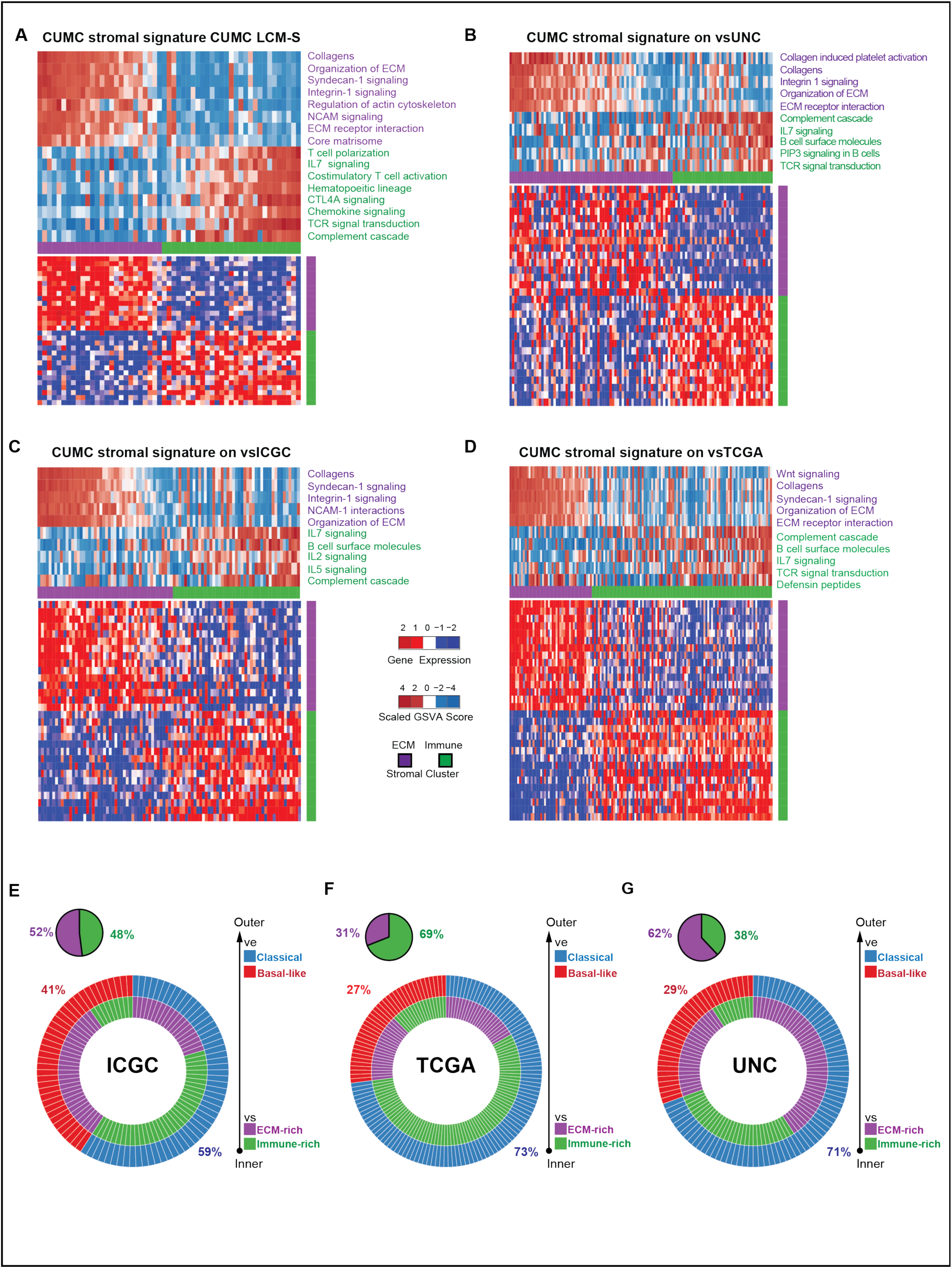
Stromal Subtyping of deconvolved external cohorts. (**A-D**) Heat-maps of the top 30 DEG between groups obtained by clustering stromal LCM–RNA–Seq samples from CUMC tumors (A), and virtual stromal (vs) profiles from the UNC (B), ICGC (C) and TCGA (D) cohorts, respectively. Clustering was based on the expression of signature derived from stromal LCM profiles from 110 individual patients (CUMC-S classifier, see Supplementary Methods). Top section of heat-map depicts GSVA scores per sample for indicated gene sets. In each virtual stroma data set, two groups were identified, one with features indicating elevated extracellular matrix deposition and remodeling (“ECM–rich”, purple) and another enriched in various immune and interleukin pathways (“Immune-rich”, green). (**E-G**) Multilayered donut plots showing (i) the alignment of epithelial with stromal subtypes for each tumor in each cohort and (ii) the proportion of each epithelial subtype. Separate pie charts summarize the proportion of stromal subtypes per cohort.

### Epithelial and stromal subtypes are partially linked and associated with survival differences

Having determined the epithelial and stromal subtypes of all CUMC, UNC, ICGC, and TCGA samples, we were able to assess the degree of variation in the subtype composition of these datasets. Notably, we found that both the epithelial and stromal composition of each tumor cohort varied considerably across datasets. Within the epithelium, the Basal–like group comprised 29%, 41%, and 27% of cases in the veUNC, veICGC, and veTCGA cohorts, respectively (Figures 5 E–G), and 36% of our epithelial LCM–RNA–Seq profiles (Figure S6A). Within the stroma, the ECM–rich subtype comprised 62%, 52%, and 31% of cases in the vsUNC, vsICGC, and vsTCGA cohorts, respectively (Figure 6 D-F), and 47% of our stromal LCM–RNA–Seq samples (Figure S6A). These observations serve to further highlight the significant heterogeneity between independent collections of pancreatic tumor specimens.

**Figure 6.**
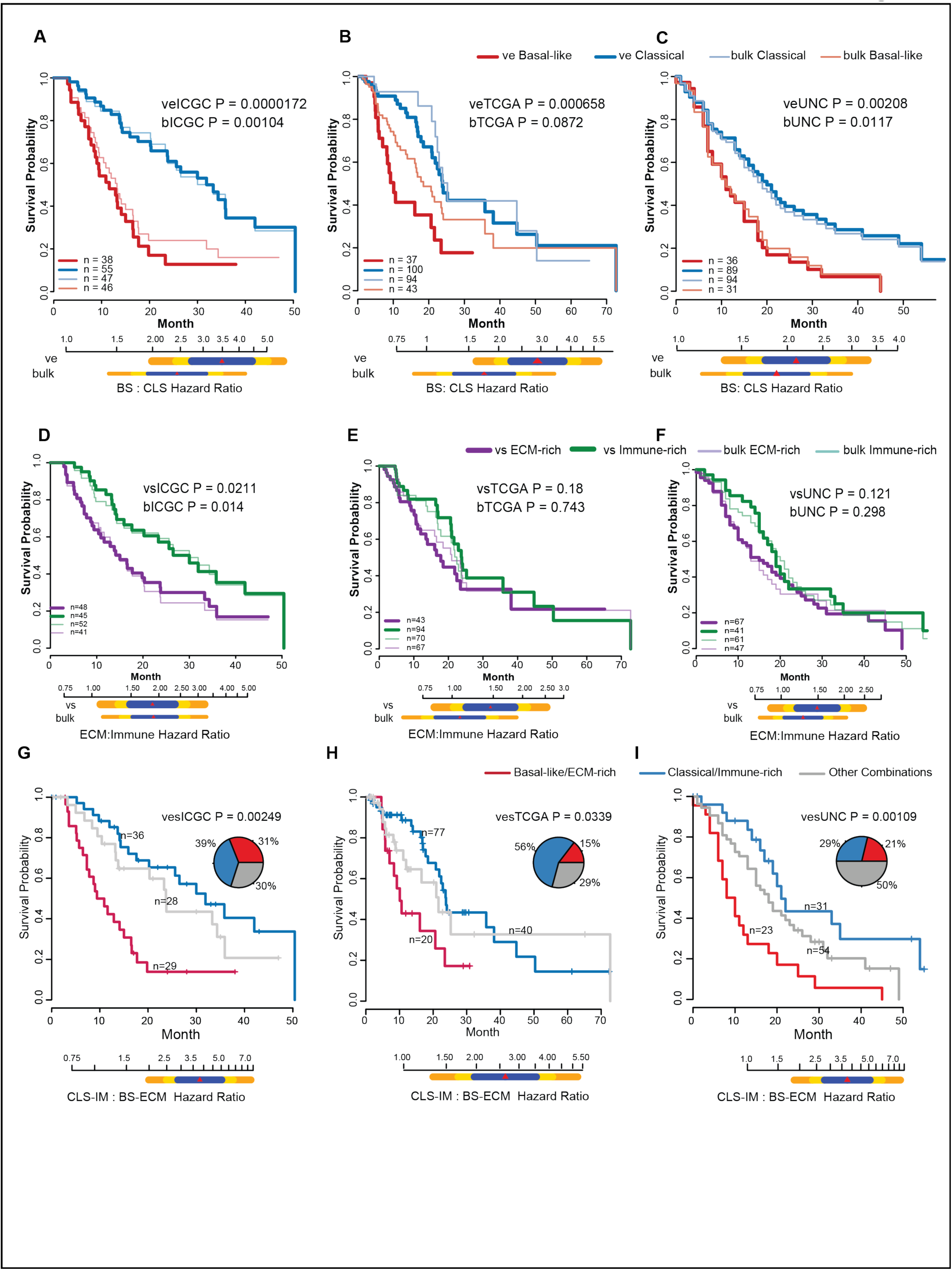
Combined epithelial and stromal subtypes associate with overall survival. (**A-C**) Kaplan-Meier (KM) survival analysis of patients with resected PDA from the indicated cohorts shows that the detection of a differential prognosis among the epithelial subtypes is enhanced by transcriptional deconvolution across all cohorts. Thick red and blue lines show the survival curves of tumor groups identified among deconvolved virtual epithelium profiles, while thinner light red and blue lines represent those groups detected by subtyping bulk expression profiles. Below each KM-plot, hazard ratios (HR) from a Cox proportional hazards model (CPHM) and their 80% (blue), 90% (yellow) and 95% (orange) confidence interval are compared between Basal-like and Classical tumors as detected in virtual epithelial (ve) and bulk expression profiles, respectively. (**D-F**) Kaplan-Meier (KM) survival analysis depicts overall survival relative to stromal subtype. Classifications derived from bulk tumor profiles are shown with thin lines while those derived from virtual stroma profiles are shown with thick lines. Below each KM-plot, hazard ratios (HR) from a univariate Cox proportional hazards model (CPHM) are shown with 80% (blue), 90% (yellow) and 95% (orange) confidence intervals (CI), comparing ECM–rich and Immune–rich tumors as detected in virtual stromal (vs) and bulk tumor profiles. Stromal subtypes are only associated with outcome in the ICGC cohort with ECM-rich tumors having a worse prognosis. (**G-I**) KM survival analysis of combined epithelial and stromal subtypes in the indicated cohorts. Red lines indicate Basal-like tumors with an ECM-rich stroma while blue lines indicate Classical tumors with an Immune-rich stroma. All other tumors are represented as a grey line. Pie charts summarize the proportion of each category per cohort. HRs from a CPHM demonstrate that the combined Basal-like/ECM-rich subtype is strongly associated with reduced survival in PDA patients in all three cohorts.

We next sought to assess the associations of epithelial and stromal subtypes with survival outcomes. Examining the epithelial samples, we found that removing stromal gene expression with ADVOCATE improved the survival association between Classical and Basal–like tumors in all three bulk datasets, with a particularly strong effect on TCGA outcomes (Figures 6 A–C) where 45% of the samples were re-classified after deconvolution. For the stromal subtypes, we observed at least a trend towards reduced survival among ECM-rich tumors in all three datasets (a finding made more apparent by deconvolution); however, this only reached significance in the ICGC cohort (Figures 6D–F). Together, these data indicate that (i) differences in tumor composition between different large-scale gene expression datasets can affect the predictive power of established classifier signatures for PDA, and (ii) transcriptional deconvolution can help overcome this hurdle, improving the reproducibility of outcome prediction.

The existence of numerous paracrine signaling pathways whose activity is affected by oncogenic mutations implies that stromal transcriptional programs should be heavily influenced by epithelial identity (Laklai et al., 2016). We examined this corollary by ascertaining the association of epithelial and stromal subtypes in our experimental LCM dataset as well as in those from the virtual UNC, ICGC, and TCGA datasets. We found that in the ICGC and TCGA cohorts, the ECM-rich stroma subtype was preferentially associated with the Basal-like epithelial subtype; the UNC and CUMC cohorts trended in this direction but did not reach significance. However, a meta-analysis of the 393 samples from all four datasets yielded an Odds Ratio of 2.7 for the association of Basal-like epithelium and ECM-rich stroma (Figure S6B, random effects model: OR 2.7 [1.33 – 5.53], p < 0.001), indicating a partial association between epithelial and stromal compartments.

The imperfect alignment of the epithelial and stromal subtypes offered the possibility that combination subtypes might vary in their survival associations as compared to either compartment alone. Indeed, consideration of combined epithelial and stromal combination subtypes affected the outcome prediction, particularly in the case of the UNC cohort where combination subtyping of deconvolved samples found a particularly poor outcome for Basal–like/ECM–rich tumors relative to Classical/Immune–rich tumors (HR = 3.76 for combined subtyping vs. 2.11 for epithelial subtyping alone (Figures 6 G–I, Figures S6 G–H). Together, these data highlight the relationship between Basal-like epithelium with ECM-rich stroma in pancreatic cancer and the strong association of this combination with poor overall survival.

## Discussion

The traditional understanding of genetic mutations as drivers of tumor development has led to a focus on the malignant compartment that is exemplified by the term “tumor purity”, which regards the stroma as mere contamination. However, with the understanding that stromal cells play critical roles in both promoting and restraining pancreatic tumor progression (Neesse et al., 2015), the consensus view of the stromal compartment has shifted to that of a critical partner – or foil – to the malignant epithelium. Indeed, in some contexts the stroma can even play a dominant role, as epitomized by the success of stroma–targeted immunotherapy in treating aggressive cancers such as metastatic melanoma and non–small cell lung cancer. In this light, we sought to study the interplay of PDA epithelium and stroma in their native state, separated by LCM from otherwise intact samples, but matched by patient so that the reciprocal signals active in each compartment might be examined.

A key outcome of this work is to unify our understanding of molecular subtypes in pancreatic ductal adenocarcinoma. To do this, we first examined the properties of subtypes resulting from existing classification schemes, making use of compartment–fraction estimation function of ADVOCATE as well as our annotation of the expression levels and compartment–specificity of each gene in our LCM-RNA-Seq dataset. We noted the substantial heterogeneity of compartment fraction between the UNC, ICGC, and TCGA cohorts, and detailed how certain proposed subtypes may have emerged from variations in tissue composition. Indeed, the removal of stromal expression signals from the Moffitt–E signature resulted in the reclassification of nearly half the samples and improved discernment of the functional processes associated with the Classical and Basal–like subtypes in each cohort.

Functional similarity in subtypes is an important concept for tumor classification. It is fully possible, even commonplace, to generate functionally distinct groups when applying the same classification signature to different sample cohorts. Variation in technical features such as library preparation method or expression platform, as well as biological heterogeneity can have an outsized impact on sample classification systems. Thus, in our effort to establish a novel classification system for PDA stroma, we placed the greatest emphasis on the reproducibility of molecular phenotypes across multiple cohorts. Following this process, we observed with great interest the emergence of two prominent molecular subtypes in the stroma with pronounced enrichment for two different aspects of stromal biology: ECM deposition and remodeling versus immune–related processes. The dichotomy between these two functional groups emphasizes the now well–described role that the inflammatory microenvironment plays in modulating the local immune response to PDA.

Examining both the epithelial and stromal subtypes together in combination across 393 pancreatic tumor specimens led to the finding that there is considerable heterogeneity in cross-compartmental dependencies across the 393 specimens examined, with some evidence of dependence between the global stromal transcriptional program and its epithelial counterpart. An expansive body of literature has accumulated describing myriad signaling interactions between the epithelium and stroma of pancreatic cancer (Bailey and Leach, 2012). Furthermore, mutation–driven epithelial signaling programs such as those induced by oncogenic K–ras have profound impacts on stromal cell biology (Pylayeva-Gupta et al., 2012; Ying et al., 2012). However, K–ras itself is mutated in 95% of PDA cases, so it cannot alone explain the differences between the prominent stromal subtypes. By examining all four tumor cohorts, we found a strong association between an ECM–rich stroma and Basal-like epithelium. The latter finding corroborates the concept that epithelial traits promoting dedifferentiation in PDA, such as the loss of SMAD4 expression, may in fact shape a more matricellular and likely more, rigid stromal phenotype (Laklai et al., 2016). This association was less prominent in the UNC cohort and absent in our smaller group of LCM tumors. Combined evidence from all cohorts, however, generally supports the idea of a modest cross-compartment dependence. Further studies will be needed to better understand the dynamics of cross-compartment subtypes in PDA by taking into account additional variables such as mutation status or environmental/epidemiological factors. To this end, we provide two novel tools for the PDA field: a compartment-specific gene signature that may discriminate between ECM-rich and Immune-rich stromas as well as the ADVOCATE framework which may help to assess the compartment composition of PDA bulk expression profiles and provide virtually purified expression for downstream analyses.

It is noteworthy that patient stratification according to both the epithelial and stromal combination subtype is strongly associated with patient outcome. Specifically, tumors with Basal–like epithelium and an ECM–rich stroma have a substantially worse overall survival compared to those with a Classical epithelium and Immune–rich stroma (HR = 3.76, 3.81, and 2.63 for UNC, ICGC, and TCGA, respectively). This effect size compares favorably to other known single variables in pancreatic cancer biology, including lymph node status (HR = 1.5), postoperative CA19-9 level (HR = 3.6) or the number of high penetrance driver genes (HR = 1.4)(Berger et al., 2008; Yachida et al., 2012) while also providing a biological context. Unfortunately, differences in the clinicopathological data reported for each cohort precluded a more sophisticated mutivariate model. Nonetheless, we expect that this approach to subtyping will have immediate applications, for example, in interpreting the results of small-scale clinical trials where random inequalities of molecular subtypes could dramatically affect the expected survival between groups or relative to historical controls.

The generation of virtual compartment–specific expression profiles from bulk profiles is a unique tool in the computational biology field. Other algorithms often provide a means to infer compartment expression from a large group of samples based on linear modeling of putative marker gene expression (Gill et al., 2014; Kuhn et al., 2011), which depends on the availability of truly cell-type specific marker genes being expressed in an essentially binary manner. By contrast, ADVOCATE uses a machine learning approach to model the expression distribution for hundreds of genes discriminating between the training conditions (i.e. epithelium and stroma). This both informs its compartment fraction prediction function and enables the deconvolution function to be performed on an individual–sample basis (provided appropriate reference standards for the sample are available from which scaled normalized expression may be obtained). The ADVOCATE framework may easily be expanded to include additional compartments or cell types, given suitable experimental data. However, it cannot generate virtual profiles for any compartments that have not been sampled; it can only estimate the fraction of residual gene expression that is unaccounted for by the modeled compartments. In practice, this “Residuals” compartment is influenced both by technical variations such as expression platform as well as biological variation such as the inclusion of non-sampled cell types. It is therefore useful as a noise-reduction tool as exemplified in Figure 4A. Considering the technical challenges of performing LCM–RNA–Seq and other enrichment methods, we expect this computational approach will reduce the need for experimental sample manipulation in the future. Moreover, while we presented implementations of ADVOCATE specific to pancreatic and breast cancer, we expect that the algorithm, which requires very limited sample sizes for training, can be generally applied to model subpopulations across the cancer field. This will provide an important framework for handling the cellular heterogeneity of cancer and further expanding the utility of large–scale gene expression profile collections.

## Experimental procedures

The information provided here is a succinct summary of the experimental procedures. Extensive and detailed information is provided in supplementary information.

### Samples studied

Information is provided from a total of 129 PDA patients who underwent surgery at the Columbia Pancreas Center. Of these, LCM–RNA–Seq data are presented for both the epithelium and stroma of 64 cases, for the stroma of an additional 59 cases, bulk RNA–Seq data are presented for 9 cases, and both LCM and bulk RNA–Seq are presented for 6 cases. Patients provided surgical informed consent which was approved by a local ethics committee (IRB # AAAB2667). Samples were frozen intraoperatively by the Columbia University Tumor Bank.

### Laser capture microdissection and RNA sequencing

Cryosections of OCT–embedded tissue blocks were transferred to PEN membrane glass slides and stained with cresyl violet acetate. Adjacent sections were H&E stained for pathology review. Laser capture microdissection was performed on a PALM MicroBeam microscope (Zeiss), collecting at least 1000 cells per compartment. RNA was extracted and libraries prepared using the Ovation RNA-Seq System V2 kit (NuGEN). Libraries were sequenced to a depth of 30 million, 100bp, single-end reads.

### Computational methods

The ADVOCATE framework is described in detail in supplementary materials. An R package for implementing ADVOCATE has been developed and will be published in advance of publication.

### Accession numbers

Expression data were deposited into publically accessible databases with accession numbers to be provided prior to publication.

## Author Contributions

H.C.M. and S.R.H. developed experimental methodologies and performed LCM–RNA–Seq. J.H. and M.B. developed the ADVOCATE algorithm and wrote related software. J.Z. provided guidance on deconvolution methods. J.H. and H.C.M. curated datasets. J.H., H.C.M., M.B., and K.P.O. analyzed and interpreted results. J.A.C., H.H., A.A, and T.S. contributed to tissue procurement and banking, and A.A. performed preliminary pathology assessments of banked samples. A.C.I. and A.R.S. marked regions of each sample to be laser capture microdissected on adjacent sections. A.C.I. annotated the histopathological features of each sample with further review of select cases by K.P.O. Regulatory oversight for the use of human samples/data was provided by H.H. and J.G. The manuscript was written by J.H., H.C.M., and K.P.O. with editing by A.C., M.B., A.C.I., and J.G. Figures were constructed by J.H., H.C.M., and K.P.O. Supervision and oversight were provided by A.C., M.B., and K.P.O.. Project was initially conceived, funded, and administered by K.P.O..

## Acknowledgements

The authors would like to thank Federico M. Giorgi and Alexander Lachmann for advice on computational methods, and Richard Moffitt for valuable critique of the manuscript. This work was supported by the National Cancer Institute (NCI) Cancer Target Discovery and Development program (1U01CA168426 to A.C.), NCI Research Centers for Cancer Systems Biology Consortium (1U54CA209997 to A.C. and K.P.O.), NCI Outstanding Investigator Award (R35CA197745-02 to A.C.) and NCI Research Project Grant (R01CA157980 to K.P.O.). Financial support was also provided by the Columbia University Pancreas Center. H.C.M. received support from a Mildred Scheel Postdoctoral Fellowship (Deutsche Krebshilfe). P.E.O. received support from the NIH NCATS (KL2TR001874).

