## Supplementary Materials for "Transcriptional deconvolution reveals consistent functional subtypes of pancreatic cancer epithelium and stroma"

### Sample acquisition

Freshly frozen tissue samples of pancreatic ductal adenocarcinoma (PDA) (n = 129) were obtained from patients who underwent surgical resection at the Pancreas Center at Columbia University Medical Center. The clinical data of these patients are shown in Supplementary Tables 1 and 2. Prior to surgery, all patients had given surgical informed consent, which was approved by institutional review board. Immediately after surgical removal, the specimens were cryopreserved, sectioned and microscopically evaluated by the Columbia University Tumor Bank (IRB AAAB2667). Suitable samples were transferred into OCT medium (Tissue Tek) and snap frozen in a 2-methylbutane dry ice slurry. The tissue blocks were stored at $-80°C$ until further processing. H&E stained sections of frozen PDA samples from the Tumor Bank were initially screened to confirm diagnosis and overall sample RNA quality was assessed by the Pancreas Center supported Next Generation Tumor Banking program using gel electrophoresis, with samples exhibiting high RNA quality utilized for subsequent analyses

### Sample extraction

Frozen tissue specimens were cut at 8 - 9 $\mu$m thickness and 2 - 3 sections were transferred onto a PEN membrane glass slide (Arcturus, Applied Biosystems). For initial histopathological review, immediate-adjacent sections were cut and stained using a standard H&E protocol to confirm the diagnosis and identify suitable areas for either laser capture microdissection or macrodissection.

### Laser capture microdissection

Throughout the staining procedure, RNAse-free water was used. Sections were fixed in 95% ethanol and stained with cresyl violet acetate (1% in Trisbuffered 70% ethanol), followed by a brief washing step in 70% ethanol, and a final dehydration in 100% ethanol. Laser capture microdissection was performed on a PALM MicroBeam microscope (Zeiss) to collect at least 1000 cells per compartment. Samples were microdissected from regions of frank carcinoma marked in advance by a GI Pathologist (A.C.I.). Captured cells were then transferred to RLT plus buffer (Qiagen) and lysed for 30 min at room temperature (RT).

### Macrodissection

For bulk tumor sections, areas containing predominantly normal or atrophic pancreas, lymphatic aggregates, larger blood vessels, or nerves were grossly trimmed from the frozen section on a PEN membrane glass slide using a sterile scalpel. Sections were then transferred to RLT plus buffer (Qiagen) and lysed for 30 min at RT.

### RNA

RNA was extracted using the RNeasy Plus Micro Kit (Qiagen) following the manufacturer's instructions. Prior to further processing, RNA integrity and yield were determined using an Agilent 2100 Bioanalyzer (RNA 6000 Pico Kit for LCM and RNA 6000 Nano Kit for bulk samples, respectively). Yields ranged from 1 to 10 ng per LCM sample and several µg per bulk sample, respectively. Only samples with a RNA Integrity Number (RIN) of at least 7 were used for further processing.

### RNA amplification and library preparation

For microdissected samples, 1 - 2 ng of RNA from LCM samples were amplified using the Ovation RNA-Seq System V2 Kit (NuGEN) following the manufacturer's instructions. The resulting cDNA libraries were fragmented using a Covaris S2 Sonicator. For samples that underwent macrodissection, a minimum of 200 ng of total RNA underwent a poly-A pull-down to enrich for mRNAs which then were used as input for the Illumina TruSeq RNA prep kit. Both types of samples were prepared for the Illumina HiSeq 2000 platform using a Beckmann-Coulter Roboter and the SPRIworks Fragment Library Kit I. Finally, a PCR using the KAPA PCR Amplification Kit was carried out. The libraries were then sequenced by the Columbia Genome Center to generate 30 million single-end reads of 100 bp length.

### RNA-Seq analysis

Reads were mapped to the human reference genome (NCBI/build 37.2) using Tophat (Version 2.0.4)(1) and two methods of gene expression quantification were employed with standard settings: (i) HTSeq (2) to obtain raw read counts per gene and (ii) Cufflinks (3) (version 2.0.2) to obtain fragments per kilobase of exon per million reads mapped (FPKM) per gene and transcript, respectively. In addition to the NCBI RefSeq gene model, gene expression was quantified using Ensembl GRCh37 gene annotations and the pipeline described, to allow the accurate evaluation of subtype-specific genes as described in Bailey et al.(4). The RSeQC package (5) was used to evaluate the suitability of RNA-Seq libraries for further analysis.

### ADVOCATE algorithm

#### Approach

The foundation of the ADVOCATE algorithm is the use of a Machine Learning based boosting algorithm to combine weak evidence derived from the expression of individual genes into a model that provides an optimal estimate of the compartment-specific composition of a heterogeneous bulk tissue. The approach assumes that the gene expression probability density function

(PDF) of the bulk is modeled as a mixture of two distinct PDFs, representing the stromal and epithelial compartments, with the optional inclusion of a third "*residual*" or "*unspecified*" compartment. The latter can be used to model either infiltration by an unknown tissue type in a specific sample or the contribution of a platform-specific bias. Inclusion of the additional compartment allows effectively addressing variability that is not statistically independent across all genes (i.e., uncorrelated noise), thus improving prediction of epithelial and stromal specific compartment representation.

**Notations**

- $x_b,\ x_e,\ x_s$: bulk, epithelial and stromal gene expression.
- $\mathcal{D}_e(\cdot)$, $\mathcal{D}_s(\cdot)$: Gaussian probability density function (PDF) of the epithelial and stromal compartments
- $\vec{p}, p_e, p_s, p_r$: gene-wise fractions. The first is a vector representing the epithelial, stromal, and residual (i.e., Residual compartment) fraction of the expression of one gene, with length of the compartment number; the second to fourth variables represent the epithelial, stromal, and residual compartment fraction, respectively.
- $\mu_e,\ \mu_s,\ \sigma_e,\ \sigma_s,\ \sigma$: the first two variables are sample mean expressions of one gene for epithelial and stromal compartment, respectively. The third, fourth and last are standard deviations for the epithelial, stromal compartment, and global, respectively.
- $\Phi(\cdot)$: Gaussian Cumulative Density Function (CDF) from training set analysis.
- $\epsilon$: user-specified prediction accuracy parameter. A smaller $\epsilon$ represents a higher prediction accuracy. For example, $\epsilon = 0.1$ represents a maximum of error 10% for sample fraction prediction.

- $k$: number of compartments, i.e. $k = 2$ or $k = 3$
- $C$: solution space defined by $\frac{1}{\epsilon}$ and $k$.
- $\vec{\omega}$, $\omega_i$: weights vector for all solutions ($C$) given a prediction accuracy ($\epsilon$) and number of compartments (k). $\vec{\omega_0}$ is the initial uniform distribution, representing that all solutions have equal chances of being the optimal solution. $\omega_{g,i}$ represents weight for the $i^{th}$ solution in $C$ after $g$ iterations.
- $l_j^g$: loss function of $j$-th solution at $g^{th}$ iteration.
- $\Delta$: set of $n$ differentially expressed genes.
- $\beta$: learning rates, a vector of size $n$. It is parameterized by the p-value derived from the negative binomial test between the epithelial and stromal LCM training samples. Each gene has a different $\beta$. $\beta_g = 1$ represents no learning. $\beta_g = 0.5$ represent the maximum learning rate, which is the learning rate of the top $\Delta_g$.
- $\tau$: threshold used in the three component model. If both $x_b$ is out of the range defined by $[\Phi_-^{-1}(\tau),\ \Phi_+^{-1}(\tau)]$, then $p_r$ has a non-zero value.
- $\vec{c}$: a vector of length k with norm = 1. So $c = [c_e, c_s]$ for k = 2.

#### Data processing

Raw counts from LCM RNA-Seq were normalized using the variance stabilizing transformation (VST) from the 'DESeq2' (6) R (7) package after filtering out genes with fewer than 5 reads in 50% of samples, with the setting: *varianceStabilizingTransformation(obj, blind = FALSE)*. Z-score transformation was then used to scale gene expression across all samples.

#### Learning

A Gaussian mixture model is used to learn the probability density function for each gene in each compartment.

$$\mathcal{D}_e(\cdot) = \sum_{i=1}^{2} \pi_{ei}\mathcal{D}(\mu_{ei}, \sigma_e^2), \quad i = \text{argmax}(\pi_{ei}) \tag{1a}$$

$$\mathcal{D}_s(\cdot) = \sum_{i=1}^{2} \pi_{si}\mathcal{D}(\mu_{si}, \sigma_s^2), \quad i = \text{argmax}(\pi_{si}) \tag{1b}$$

$$\sigma = \sqrt{\sigma_e^2 \frac{n_e - 1}{n_e + n_s} + \sigma_s^2 \frac{n_s - 1}{n_e + n_s}} \tag{1c}$$

#### 2-component model gene-specific fraction prediction

In a 2-component model, the bulk expression PDF of a gene is assumed to be the weighed sum of its epithelial and stromal PDFs only.

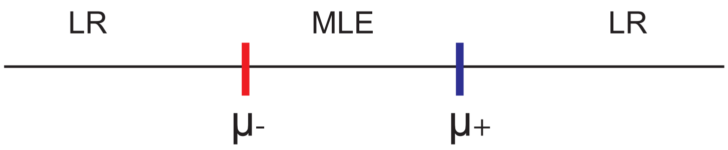


2-component model

**Maximal Likelihood (MLE)** The objective function here is to identify the parameters so that the probability of observing a given compartment-specific distribution for each gene is maximized.

$$\text{argmax}(\ \mathcal{D}_e(X_e = x_e) \cdot \mathcal{D}_s(X_s = x_s)\ ) \tag{2}$$

$$\begin{cases} p_e x_e + p_s x_s = x_b \\ p_e + p_s = 1 \\ p_e \geq 0; p_s \geq 0 \end{cases} \tag{3}$$

$$X_e = \mathcal{D}(\mu_e, \sigma)\ ; X_s = \mathcal{D}(\mu_s, \sigma) \tag{4}$$

Rewriting object function (Equation 2) using Equation 1, we have:

$$\text{argmax}(\ \mathcal{D}_e(X_e = x_e) \wedge \mathcal{D}_s(X_s = x_s)) \quad \rightarrow \tag{5}$$

$$\text{argmin}\ (\ \log\ \mathcal{D}_e(X_e = x_e) + \log\ \mathcal{D}_s(X_s = x_s)\ ) \quad \rightarrow \tag{6}$$

$$\text{argmin}\ (log(\frac{1}{\sqrt{2\pi\sigma^2}}exp(\frac{x_e - \mu_e}{2\sigma^2})^2) + log(\frac{1}{\sqrt{2\pi\sigma^2}}exp(\frac{x_s - \mu_s}{2\sigma^2})^2)) \tag{7}$$

In this context, we define:

$$\mu_- = \min(\mu_e, \mu_s)$$
$$\mu_+ = \max(\mu_e, \mu_s)$$

Based on Maximum Likelihood Estimation (MLE), the above equation reaches a minimum for $x_e = \mu_e$ and $x_s = \mu_s$ (7). Thus, substituing in Equation (3), one has:

$$p_e \cdot \mu_e + (1 - p_e) \cdot \mu_s = x_b \tag{8}$$

$$p_e \cdot (\mu_e - \mu_s) = x_b - \mu_s \tag{9}$$

$$p_e^{\text{MLE}} = \frac{x_b - \mu_s}{\mu_e - \mu_s} \tag{10}$$

**Likelihood Ratio (LR)** It is easy to observe that in Equation (10), $p_e$ is only defined positive for $x_b \in (\mu_-,\ \mu_+)$.

When $x_b \leq \mu_-$, we use the likelihood ratio between $\mathcal{Q}_e = \mathcal{D}_e(X_e = x_b)$ and $\mathcal{Q}_s = \mathcal{D}_s(X_s = x_b)$ to compute $p_e$. Here, $\mathcal{Q}_e, \mathcal{Q}_s$ represent likelihood before normalization.

$$\mathcal{Q}_e = \Phi_e(x_b) \tag{11}$$

$$\mathcal{Q}_s = \Phi_s(x_b) \tag{12}$$

After normalization and substitution, we have equation (13).

$$p_e = \frac{\mathcal{Q}_e}{\mathcal{Q}_e + \mathcal{Q}_s}, \text{ if } x_b \in (-\infty, \mu_-] \tag{13}$$

when $x_b \geq \mu_+$, an identical approach can be used. Thus, we have equation (14) to compute gene-specific fraction in a 2-compartment model.

$$p_e = \begin{cases} \frac{x_b - \mu_s}{\mu_e - \mu_s}, & \text{if } x_b \in (\mu_-,\ \mu_+) \\ \frac{1-\mathcal{Q}_e}{2-\mathcal{Q}_e-\mathcal{Q}_s}, & \text{if } x_b \in [\mu_+,\ \infty) \\ \frac{\mathcal{Q}_e}{\mathcal{Q}_e+\mathcal{Q}_s}, & \text{if } x_b \in (-\infty, \mu_-] \end{cases} \tag{14}$$

$p_s$ is calculated using Equation 15.

$$p_s = \ 1 - p_e \tag{15}$$

Thus, the compartment-specific contribution to the expression of gene $g$ is given by (Equation 16):

$$\vec{p} = \ [p_e,\ p_s] \tag{16}$$

**3-component model gene-specific fraction prediction**

In a three-component model, the bulk PDF of a gene is computed as a mixture of its epithelial PDF, its stromal PDF and the PDF of an additional "residual" or "unspecified" compartment. The latter is used to account for correlated variability across genes due to sample-specific infiltration of unknown tissue or introduction of platform-specific bias.
Use of a 3-compartment model is formulated by specifying a threshold $\tau$ to determine a range of bulk gene expression ($[x_{\text{-}}(\tau),\ x_+(\tau)]$) that can be effectively modeled using only the epithelial and stromal PDFs, i.e., where the probability of bulk expression is greater than $\tau$ (e.g., $\tau = 10^{-5}$).

$$\text{argmax}(\ \mathcal{D}_e(X_e = x_e) \cdot \mathcal{D}_s(X_s = x_s) \cdot \mathcal{D}_r(\cdot)\ ) \tag{17}$$

**Algorithm 1:** Gene-specific compartmental fraction inference

**Input:** $x_b$, $D_e(\mu_e, \sigma^2)$, $D_s(\mu_s, \sigma^2)$

**Output:** $\vec{p}$

**if** $\mu_- < x_b < \mu_+$ **then**

  $p_e^{\mathrm{MLE}} = \frac{x_b - \mu_s}{\mu_e - \mu_s}$

$\mathcal{Q}_e = \Phi_e(x_b); \quad \mathcal{Q}_s = \Phi_s(x_b)$

**if** $x_b \leq \mu_- < \mu_+$ **then**

  $p_e = \frac{\mathcal{Q}_e}{\mathcal{Q}_e + \mathcal{Q}_s}$

**if** $x_b \geq \mu_+ > \mu_-$ **then**

  $p_e = \frac{1 - \mathcal{Q}_e}{2 - \mathcal{Q}_e - \mathcal{Q}_s}$

$p_s = 1 - p_e$

**return** $\vec{p} = [p_e,\ p_s]$.

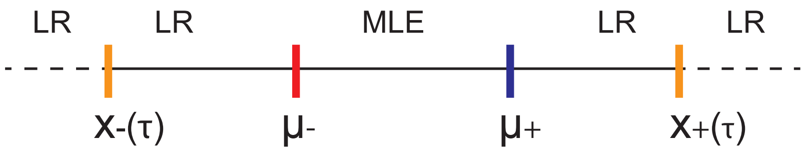


3-component model

$$\begin{cases} p_e x_e + p_s x_s + p_r x_r = x_b \\ p_e + p_s + p_r = 1 \\ p_e \geq 0, p_s \geq 0, p_r \geq 0 \end{cases} \tag{18}$$

$$X_e = \mathcal{D}(\mu_e, \sigma); X_s = \mathcal{D}(\mu_s, \sigma) \tag{19}$$

**MLE and LR** when $x_b \in [x_-(\tau),\ x_+(\tau)]$, Equation (14) still applies, with $p_r = 0$.

$$x_-(\tau) = \min(\Phi_e^{-1}(\tau), \Phi_s^{-1}(\tau)); \tag{20}$$
$$x_+(\tau) = \max(\Phi_e^{-1}(\tau), \Phi_s^{-1}(\tau)) \tag{21}$$
$$\tag{22}$$

**Adjusted Likelihood** when $x_b < x_-(\tau) < \mu_-$ or $x_b > x_+(\tau) > \mu_+$, we define $p_r = 1 - \mathcal{Q}_e - \mathcal{Q}_s$, with $\mathcal{Q}_e$ and $\mathcal{Q}_s$ representing the epithelial and stromal fractional composition likelihoods before normalization. Thus, Equation 24 follows.

$$\mathcal{Q}_e = \mathcal{D}_e(x_b); \quad \mathcal{Q}_s = \mathcal{D}_s(x_b) \tag{23}$$

$$p_r = \begin{cases} 1 - \mathcal{Q}_e - \mathcal{Q}_s, & x_b \in (-\infty, x_-(\tau)) \cup (x_+(\tau), \infty) \\ 0, & \text{Else} \end{cases} \tag{24}$$

Taking together, in a 3-component model, one can compute $p_e$ and $p_s$ as :

$$p_e = \begin{cases} \frac{x_b - \mu_s}{\mu_e - \mu_s}, & \text{if } x_b \in (\mu_-, \mu_+) \\ \frac{\mathcal{Q}_e}{\mathcal{Q}_e + \mathcal{Q}_s}, & \text{if } x_b \in [x_{-\tau}, \mu_-] \\ \frac{1 - \mathcal{Q}_e}{2 - \mathcal{Q}_e - \mathcal{Q}_s}, & \text{if } x_b \in [\mu_+, x_+(\tau)] \\ \mathcal{Q}_e, & \text{if } x_b \in (-\infty, x_+(\tau)) \\ 1 - \mathcal{Q}_e, & \text{if } x_b \in (x_-(\tau), \infty) \end{cases} \tag{25}$$

$$p_s = \begin{cases} \frac{x_b - \mu_-}{\mu_+ - \mu_-}, & \text{if } x_b \in (\mu_-,\ \mu_+) \\ \frac{\mathcal{Q}_s}{\mathcal{Q}_e + \mathcal{Q}_s}, & \text{if } x_b \in [x_-(\tau), \mu_-] \\ \frac{1 - \mathcal{Q}_s}{2 - \mathcal{Q}_e - \mathcal{Q}_s}, & \text{if } x_b \in [\mu_+, x_+(\tau)] \\ \mathcal{Q}_s, & \text{if } x_b \in (-\infty, x_-(\tau)) \\ 1 - \mathcal{Q}_s, & \text{if } x_b \in (x_+(\tau), \infty) \end{cases} \tag{26}$$

This allows estimating the compartment-specific fractional composition $\overrightarrow{p}$, as in equation 27.

$$\overrightarrow{p} = [p_e,\ p_s,\ p_r] \tag{27}$$

**Algorithm 2:** Gene-specific compartmental fraction inference

**Input:** $x_b$, $\mathcal{D}_e(\mu_e, \sigma^2)$, $\mathcal{D}_s = (\mu_s, \sigma^2)$,$\tau$

**Output:** $\overrightarrow{p} = [p_e,\ p_s,\ p_r]$

**if** $\mu_s < x_b < \mu_e$ **then**

- $p_e^{MLE} = \frac{x_b - \mu_s}{\mu_e - \mu_s}$
- $p_s^{MLE} = \frac{x_b - \mu_e}{\mu_s - \mu_e}$
- $p_r = 0$

$\mathcal{Q}_e = \Phi_e(x_b); \quad \mathcal{Q}_s = \Phi_s(x_b)$

$x_-(\tau) = \min(\Phi_e^{-1}(\tau), \Phi_s^{-1}(\tau))$;

$x_+(\tau) = \max(\Phi_e^{-1}(\tau), \Phi_s^{-1}(\tau))$

**if** $x_-(\tau) \leq x_b \leq \mu_-$ **then**

- $p_e = \frac{\mathcal{Q}_e}{\mathcal{Q}_e + \mathcal{Q}_s}$
- $p_s = \frac{\mathcal{Q}_s}{\mathcal{Q}_e + \mathcal{Q}_s}$
- $p_r = 0$

**if** $x_+(\tau) \geq x_b \geq \mu_+$ **then**

- $p_e = \frac{1 - \mathcal{Q}_e}{2 - \mathcal{Q}_e - \mathcal{Q}_s}$
- $p_s = \frac{1 - \mathcal{Q}_s}{2 - \mathcal{Q}_e - \mathcal{Q}_s}$
- $p_r = 0$

**if** $x_b < x_-(\tau)$ **then**

- $p_e = \mathcal{Q}_e$
- $p_s = \mathcal{Q}_s$
- $p_r = 1 - \mathcal{Q}_e - \mathcal{Q}_s$

**if** $x_b > x_+(\tau)$ **then**

- $p_e = 1 - \mathcal{Q}_e$
- $p_s = 1 - \mathcal{Q}_s$
- $p_r = 2 - \mathcal{Q}_e - \mathcal{Q}_s$

**return** $\overrightarrow{p} = [p_e,\ p_s,\ p_r]$.

#### Sample-specific fraction prediction

Based on the machine learning theory proposed by Yoav Freund (26), a multiplicative weighted-majority procedure was designed to integrate individual weak evidence represented by compartment-specific fractional composition predicted on an individual gene basis to predict an optimal compartment-specific fractional composition for the whole sample $\vec{c}$ with a maximum, user-specified prediction error ($\epsilon$).

After running the algorithms 1 or 2, the individual gene-specific fractional composition values ($\vec{p}$) are used as an input to the final evidence integration step, as represented by algorithm 3.

ADVOCATE boosting procedure starts by creating an initial solution space ($C$) with a set of solutions such that at least one of them is guaranteed to be within an $\epsilon$ of the true solution. The dimensionality of this space is determined by $k = 2$ or $k = 3$. A set of weights $\omega_0$ representing the posterior probability of each solution in $C$ is initialized to a uniform distribution. The total number of solutions ($N_\epsilon^{k-1}$) grows as $\epsilon$ becomes smaller and with increasing values of $k$.

$$\begin{aligned}&\text{Given } \epsilon, \text{for k-component model}\\ &\mathcal{C} := [0, \epsilon, 2\epsilon, \ldots, 1]^{k-1}\\ &\vec{\omega}_0 := [\epsilon^{k-1}]\end{aligned}$$

$C^2$ represents the solution space for a 2-compartment model, when $\epsilon = 0.1$, degree of freedom = 1, thus, learning on 1-D

$$\begin{bmatrix} 0 & 0.1 & 0.2 & \ldots & 0.7 & 0.8 & 0.9 & 1 \\ 1 & 0.9 & 0.8 & \ldots & 0.3 & 0.2 & 0.1 & 0 \end{bmatrix}$$

The $\omega_0^1$ vector for $C^2$ is thus:

$$\begin{bmatrix} 0.0909 & 0.0909 & 0.0909 & \ldots & 0.0909 & 0.0909 & 0.0909 & 0.0909 \end{bmatrix}$$

$C^3$ for represents the solution space for a 3-compartment model when $\epsilon = 0.1$, degree of freedom = 2, thus learning on 2-D.

$$\begin{bmatrix} 0 & \dots & 0.1 & \dots & 0.5 & \dots & 0.9 & 1 \\ 0 & \dots & 0.1 & \dots & 0.1 & \dots & 0.1 & 0 \\ 1 & \dots & 0.8 & \dots & 0.4 & \dots & 0 & 0 \end{bmatrix}$$

The $\omega_0^2$ vector for $C^3$ is thus:

$$\begin{bmatrix} 0.015 & \dots & 0.015 & \dots & 0.015 & \dots & 0.015 & 0.015 \end{bmatrix}$$

Then, ADVOCATE iteratively updates the weight vector $\vec{\omega}_g$ for each gene based on the loss function (see Equation 29). Here $\beta$ is the learning rate, as parameterized by the p-value of the differential gene expression analysis.

$$\vec{\omega}_g = \vec{\omega}_{g-1} \cdot \beta_g^{\mathcal{L}_g} \tag{28}$$

$$\mathcal{L}_{g,i} = |\mathcal{C}_i^{k-1} - p_g^{k-1}| \tag{29}$$

$$\beta_g \propto \mathcal{F}(\text{Pvalue}_g), \;\; \beta_g \in [0.5, 1] \tag{30}$$

After all iterations are completed, the solution with the maximum weight ($\vec{\omega}_{g,i}$) is chosen as the predicted sample-specific fraction compositon ($(c)$) (Equation 31).

$$i = \text{argmax}_{i \in [1, N_\epsilon^{k-1}]} \vec{\omega}_g \tag{31}$$

$$\vec{c} = \mathcal{C}_i^k \tag{32}$$

**Algorithm 3:** Prediction of sample-specific fraction

**Input:** $\Delta$, $\vec{p}$,$\vec{\beta}$,$\epsilon$, $k$

**Output:** Sample compartmental fraction $c$

$C = \{0, \epsilon, 2\epsilon, \ldots, 1\}^{k-1}$ /*$C$ is the solution space, $c_i = i \cdot \epsilon$ is the $i$-th solution */

$\omega_0 = (\epsilon^{k-1}, \ldots, \epsilon^{k-1})$ /* Initializing weights with uniform distribution over all solutions/

$N_\epsilon^{k-1} := (\frac{1}{\epsilon})^{k-1}$ /*Number of solutions*/

**for** $g \leftarrow \Delta$ **do**

  $\vec{p}_g$ = result from algorithm 1 or 2

**for** $g \leftarrow \Delta$ **do**

  **for** $i \leq N_\epsilon^{k-1}$ **do**

    $\mathcal{L}_i^g = |C_i - \vec{p_g}|$ /*Loss function of $i$-th solution at gene $g$ */

    $\omega_i = (\omega_{g-1})_i \cdot \beta_g^{\mathcal{L}_i^g}$ /*update weights according to loss function */

  Normalizing $\vec{\omega} = (\omega_1, \ldots, \omega_{N_\epsilon^{k-1}})$ to get updated $\vec{\omega_g}$

$i = \max_i\{\omega_i\}$

**return** $\vec{c} = C_i$ /* The $i$-th solution has maximum weight. */

#### Extracting compartmental specific expression

For genes $\in \Delta$ , we have

$$x_{g,e} = x_{g,b} \cdot p_{g,e} \tag{33}$$

$$x_{g,s} = x_{g,b} \cdot p_{g,s} \tag{34}$$

For genes $\notin \Delta$, we have

$$x_{g,e} = x_{g,b} \cdot c_e \tag{35}$$

$$x_{g,s} = x_{g,b} \cdot c_s \tag{36}$$

#### Power Analysis

In order to evaluate the impact of sample size on prediction accuracy, random sub-sampling of 60 LCM pairs was carried out by varying sample sizes from 5 to 60 at an interval of 5. For each sample size except 60, 100 random samples were drawn. ADVOCATE was trained using the expression of sub-selected paired LCM samples and, subsequently, used to predict the compartment fraction of all LCM samples. The prediction error was calculated by considering all epithelial and stromal LCM samples to be 100% and 0% epithelial, respectively.

### Computational validation of compartment fraction estimation

**In silico datasets.** 60 pairs of simulated compartmental LCM expression profiles were generated by random sampling from a Gaussian distribution parameterized by each gene's mean expression in all epithelial and stromal LCM samples, respectively, as the means and each gene's global variance across compartments as the variances. Synthetic bulk expression profiles were then generated by linearly mixing simulated epithelial and stromal expression profiles at different fractions, ranging from 1% to 99% epithelial fraction with and increment of 1%. Semi-synthetic bulk samples were generated by mixing actual paired compartment-specific LCM GEP following the same mixing procedure.

**Computational validation procedure.** Compartment fractions from synthetic and semi-synthetic bulk datasets were predicted by training ADVOCATE using simulated and real LCM expression, respectively. For Leave-One-Out Cross Validation (LOOCV), ADVOCATE was trained using all but one pair of samples and tested on the left-out pair. Prediction error was calculated between predicted epithelial fractions and the original mixing proportions for synthetic and semi-synthetic data, respectively. For LOOCV, it was calculated by assuming 100% epithelial and stromal cells, respectively, in each paired sample that was tested.

#### Comparison to algorithms.

To quantify the tumor purity for tumor expression profiles (whether experimental, virtual, or synthetic), we used the R package 'estimate'(14) with 'GeneSymbol' as id mapping and 'illumina' as platform. The parameters for calculating tumor purity based on ESTIMATE scores were set to the default values (a = 0.0001467884, b = 0.6049872018). Affymetrix LCM data used in Yoshihara et al. were downloaded from the Gene Expression Omnibus (GEO) using the 'GEOquery' package(15) and accession numbers GSE14548, GSE10797, GSE9890 and GSE29156. Calculations of stromal, immune and ESTIMATE scores with consequent tumor purity estimations were carried out as described above by selecting 'affymetrix' as the platform setting, with the rest of the parameters as default.

To further evaluate algorithm performance, we compared the all predictions from ESTIMATE, PSEA (16), DSA (17), and deconRNAseq (18) with ADVOCATE. We used the marker genes (Figure 1) as the reference genes. R function *lsqnonneg* from package R 'pracma' was used to solve linear equations in PSEA, the same way as suggested by (16). R function *'solve'* from 'MASS' function was used to compute weights for DSA. Raw-counts of 60 pairs of training LCMs together with the same DEGs used in ADVOCATE training, were used to train *deconRNAseq*.

### Experimental validation

#### Assessment of RNA amplification behavior.

In order to test whether RNA amplification using the Ovation RNA-Seq System V2 (NuGEN) indeed occurs in a linear manner (22), we added ERCC

Spike-In Mix 1 (Ambion) at increasing concentrations (1X, 2X, ..., 32X - 2 replicates per concentration) to RNA samples before amplification. ERCC Spike-In Mix 1 contains a collection of 92 synthetic, polyadenylated transcripts between 250 and 2000 bp at defined concentrations. For quantification purposes, the ERCC Spike-In sequences were added to the reference genome and annotation file, respectively, and the same Tophat/HTSeq pipeline that was outlined above was used for raw read quantification. Read counts from libraries containing ERCC Spike-In mix were normalized by accounting for differences in library size using the DESeq2 package ( counts(obj, normalized = TRUE) ). ERCC Spike-In species that did not have at least one read in 80% of the spiked-in libraries were discarded from the analysis, which left 19 distinct ERCC transcripts, 16 of which could be categorized into 2 major length categories and 3 major concentration categories (see table below). Both normalized reads and Spike-In concentration were log2-transformed and Pearson correlation was determined at each level of length and concentration, respectively.

**LCM samples with bulk expression profiles.**

For cases in which both paired LCM and bulk expression profiles were available (n = 6), bulk profiles were deconvolved into virtual epithelium and stroma profiles, respectively. Pearson correlation was calculated between the actual compartment–specific LCM expression profiles and its virtually purified counterpart. Correlation between randomized bulk and LCM samples served as a control for all the other correlations. The randomized bulk expression profiles were generated by shuffling gene identifiers.

**Nuclei counting.**

15 bulk PDA sections were macrodissected as described above to enrich malignant epithelium and adjacent stroma. Sections used for RNA extraction and molecular profiling were derived immediately adjacent to sections used

Table 1: ERCC spike-ins used in LCM RNA sequencing

| **ERCC ID** | **Concentration log10(attomol/$\mu$l)** | **Concentration category** | **Length in bp** | **Length category** |
|---|---|---|---|---|
| ERCC-00002 | 4.18 | 104 - 105 | 1061 | 1000 bp |
| ERCC-00003 | 2.97 | 102 - 103 | 1023 | 1000 bp |
| ERCC-00004 | 3.88 | 103 - 104 | 523 | 500 bp |
| ERCC-00009 | 2.97 | 102 - 103 | 984 | 1000 bp |
| ERCC-00042 | 2.67 | 102 - 103 | 1023 | 1000 bp |
| ERCC-00043 | 2.67 | 102 - 103 | 1023 | 1000 bp |
| ERCC-00046 | 3.58 | 103 - 104 | 522 | 500 bp |
| ERCC-00060 | 2.36 | 102 - 103 | 523 | 500 bp |
| ERCC-00074 | 4.18 | 104 - 105 | 522 | 500 bp |
| ERCC-00095 | 2.08 | 102 - 103 | 521 | 500 bp |
| ERCC-00108 | 2.97 | 102 - 103 | 1022 | 1000 bp |
| ERCC-00111 | 2.67 | 102 - 103 | 994 | 1000 bp |
| ERCC-00130 | 4.48 | 104 - 105 | 1059 | 1000 bp |
| ERCC-00136 | 3.28 | 103 - 104 | 1033 | 1000 bp |
| ERCC-00145 | 2.97 | 102 - 103 | 1042 | 1000 bp |
| ERCC-00171 | 3.58 | 103 - 104 | 505 | 500 bp |

for histopathological assessment. For each bulk section, 8 - 10 high-power fields were recorded, epithelial and stromal areas identified and then nuclei were counted manually using the ImageJ (20) software and its built-in multi-point tool. Counts were noted per compartment and divided by the grand total to yield compartmental fractions. **M**ean **A**bsolute **P**ercentage **E**rror (MAPE) was used to quantify ADVOCATE prediction accuracy (equation (18)). The epithelial fraction based on nucleus counting or area summation was considered as validation (V). ADVOCATE's molecular prediction of epithelial fraction was considered as prediction (P).

$$\text{MAPE} = \frac{100}{n}\sum_{i=1}^{n}\left|\frac{V_i - P_i}{max(V_i, P_i)}\right|$$

**Human Protein Atlas (HPA).**

Using the HPA search algorithm, we subset the list of potential proteins to those that had highest quality antibodies for immunohistochemistry (IHC) by applying the filter 'ih_tissue_reliability:Supportive'. Next, we evaluated IHC staining patterns for PDA tumor sections in the HPA pathology database for the top and bottom 50 genes of our subset DEG list which represented mRNA-predicted stromal and epithelial genes, respectively. The observed IHC staining patterns were categorized as follows: (i) 'strongly supportive' for cases where IHC staining aligned with mRNA prediction with both high signal intensity and compartment-specificity, (ii) 'weakly supportive' for cases where IHC staining aligned with mRNA prediction with either moderate signal intensity or moderate compartment-specificity (i.e. differential expression can be appreciated on the protein level, but is not bimodal), (iii) 'indeterminate' for cases with absent compartment-specificity (i.e. no appreciable differential expression on the protein level), (iv) 'not expressed' for cases where low or absent signal intensity precluded meaningful analysis, and (v) 'opposing' for cases where IHC staining contradicted mRNA prediction. The full list of genes that are used as compartment-specific genes by ADVOCATE, the list of genes for which antibodies with high quality for IHC are available at the HPA and the results from the described analysis can be found in Supplementary Tables 3 - 5.

### ADVOCATE training and prediction on Breast Cancer Agilent array-based LCM

The breast cancer LCM epithelial and stromal array expresion data (GSE68744) was downloaded from GEO (27). Among them, 47 pairs (n=94) were used as training following the same framework used in CUMC-LCM-RNAseq PDA sample training. Leave-one-out cross validation were used for the 94 training samples. Based on this training, we did independent prediction of the remaining 25 pairs. Another validation cohort from the same publication were donwloaed from the supplementary data of (27). It includes 36 paired LCM samples. We then used ADVOCATE and ETIMATE algorithm to predict the epithelial fraction of these 36 pair LCM aligent array samples following the same setting as before.

### ADVOCATE prediction on external PDA datasets

#### Dataset preparation.

The UNC dataset was downloaded using the R package 'GEOquery'(15), and accession number GSE71729. Only pancreatic primary tumor samples with survival data were retained for our analysis (n = 125). For the ICGC cohort, normalized expression and clinical data were extracted from the supplementary information provided by Bailey et al(4). After matching sample IDs with phenotype data, we kept 93 samples for our analysis, excluding the two acinar cell carcinomas. RNA-Seq V2 data was downloaded from the TCGA data portal on 5/31/2016 together with clinical and biospecimen information

- including a report on 27 cases that were retracted after review by the PAAD EPC. In addition, we excluded cases with a diagnosis of 'other malignancy'. Raw read counts per gene were extracted from the RSEM output files for all samples, normalized to account for different library sizes, and the variance was stabilized as implemented in the DESeq2(6) package. Details on the epithelial and stromal molecular subtypes per case as well as the survival data used can be found in supplementary table 6.

**Deconvolution on external datasets.**

Given the different criteria that each of the described external studies used for including PDA samples for molecular profiling together with the substantially larger sample sizes, we expected a higher degree of tumor tissue heterogeneity and, therefore, employed the three-compartment ADVOCATE implementation (prediction accuracy ) to estimate epithelial, stromal and residual compartment fractions in these cohorts. Inferred compartment-specific virtual expression profiles were derived from the respective bulk expression profiles using the two-compartment implementation of ADVOCATE as described above. After estimating epithelial, stromal and unspecified fractions, the former two were scaled to sum up to 1. Gene signatures used for classification Clustering of both bulk expression profiles and virtually purified compartment–specific profiles was performed by sub-selecting the normalized expression data of epithelial signature genes described by Moffitt et al (21) for epithelial subtypes and genes distinguishing the two major stromal subtypes in LCM data for stromal subtypes. Both signature gene lists can be found in supplementary tables 21 and 22. Derivation of subtype-specific genes in the ICGC cohort Subtype-specific classifier genes for 'ADEX', 'Immunogenic', 'Pancreatic Progenitor' and 'Squamous' tumors were derived as follows from the list of 613 genes found to be differentially expressed between all subtypes by Bailey et al. (= SAM genes). Normalized expression data and information on subtype and silhouette score for 96 tumors from the

ICGC cohort were extracted from the supplemental data provided by Bailey et al. Only tumors with positive silhouette scores were retained for the analysis (n = 83). First, 'subtype vs. rest' comparisons were performed for each subtype using the limma R package. As an initial filtering step to identify subtype-specific genes, only those genes were kept that were significantly upregulated in the respective subtype while not being significantly upregulated in any of the other comparisons. If this included genes for which there were non-significant trends (i.e. positive t-statistics) in other comparisons, an additional comparison was made between 'subtype A and B' specifically. This strategy yielded 48 ADEX-specific, 92 Immunogenic-specific, 3 Progenitor-specific and 70 Squamous-specific genes. Input data, results from the DEG analyses and all classifier lists used in this manuscript can be found in Supplementary Tables 17-22.

### Derivation of a stromal subtype signature.

Stromal LCM-RNA-Seq gene expression profiles from 110 patients were VST-normalized as described above and subset to stroma-specific genes as determined from the paired DEG between epithelium and stroma (t-statistic > 0 and FDR < 0.1, respectively). Furthermore, genes expressed from chromosome Y as well as from the XIST (X-inactive specific transcript) locus were excluded as their variance is likely to be related to gender-specific differences in gene expression, which we reasoned should not be considered during subtyping, leaving us with 4401 genes. The 1000 most variable genes were determined from 100 bootstraps of the 4401 genes x 110 samples input matrix based on their median absolute deviation (MAD). The 1000 genes occurring most often in these 100 bootstraps were used as input for non-negative matrix factorization (NMF) using the NMF R package(10) with its standard algorithm (Brunet) and a random seeding method where entries of each factor are drawn from a uniform distribution over [0, max(A)], where A is the input matrix. The factorization rank was estimated in 50 runs of

NMF from actual and randomized data for 2 through 10 ranks. Cophenetic correlation and silhouette scores of the consensus matrix decreased substantially already from 2 to 3 ranks and declined further for higher factorization ranks. NMF was then run with 200 iterations (*nmf(x, rank = 2, nrun = 200, seed = 1)*) and the samples were assigned a cluster by hierarchical clustering of the consensus matrix using complete linkage. In order to derive a signature distinguishing between the two subtypes, differential genes were identified using the siggenes R package (11). Genes with a q-value $< 0.001$ and either an above median expression or an effect size $> 1.5$ were retained. The signature genes and their functional annotation using Gene Ontology Biological Processes and the DAVID Bioinformatics Resources (12) can be found in supplementary tables 22 - 24.

**Clustering**

A Pearson dissimilarity distance matrix was used as the input for k-means consensus clustering for each dataset with the same preset random seed value. Each clustering was run 500 times. Subsequently, the cluster output from all k-means clustering was hierarchically clustered and the number of clusters was decided based on the cutree function in R '*stats*', thereby yielding the final results for each dataset and each compartment. Using the average silhouette score, most robust results were obtained for 2 epithelial and stromal clusters, respectively, in each data set. For the virtual epithelial expression profiles in the UNC cohort, both 2 and 3 clusters were equally feasible. We elected to use 2 clusters for this dataset for the sake of consistency. In order to be consistent with the stromal signature derivation process, stromal class assignment was carried out using the same NMF and hierarchical clustering pipeline described above. As expected some CUMC stromal signature genes were less informative, i.e. showed low variance as determined by their interquartile range (IQR), in the external cohorts. Therefore, those stromal signature genes exhibiting a below global variance in a given cohort were removed before clustering.

### Gene set enrichment analysis

We used the R implementation of single sample Gene Set Enrichment analysis: GSVA (gene set variation analysis) with default parameters (22). The input expression matrix was filtered for the most variable 50% of the genes as determined by interquartile range (IQR) in case of the epithelial annotation and to stromal genes (log2 fold change $> 0.5$ in the paired epithelium vs. stroma differential expression analysis) in case of the stromal annotation. For annotation of epithelial subtypes, we tested a set of gene sets shown previously to be discriminating between classical and basal-like tumors (4,18). For the annotation of stromal subtypes, we first determined stroma-specific pathways among the c2 canonical pathways module from the MSigDB(4) by comparing the ADVOCATE epithelial and stromal training samples. Using these stroma-specific pathways (FDR in epithelial vs. stromal GSVA comparison $< 0.05$), we performed differential enrichment analysis between the prominent stromal subtypes in the LCM cohort using the R package 'limma' (23) and selected a similar number of illustrative gene sets as for the epithelium. All tested gene sets and their description can be found in supplementary tables 7 and 8.

### Silhouette score and Differential Gene Expression (DEG)

Silhouette scores were calculated using the silhouette function from the cluster (24) R package with the k-means clustering results and the Pearson dissimilarity matrix as input. Differential gene expression analysis was calculated out using the 'limma' (23) package. Among all differentially expressed genes between compartment-specific subtypes (adjusted p-value $< 0.05$) the top 30 genes per subtype were shown in the heatmaps.

### Survival analysis

The association of molecular subtypes with disease outcome was evaluated using Kaplan-Meier survival analysis calculated along with log-log p value using the surv and coxph functions from the 'survival' (25) package. Furthermore, fitting and visualization of Cox proportional hazards models were carried out using the cph function from the R package 'rms' with all ties handled using the Breslow method.

### Effect size meta-analysis.

In order to evaluate evidence from four PDA cohorts on the co-occurrence of certain compartment–specific molecular subtypes, a meta-analysis was carried out using the metafor (13) R package. The 2 x 2 table was set up in a way that Basal-like tumors and ECM-rich stromas represented the first row and column, respectively. The log odds ratio and its variance were calculated for each study using the *escalc(measure = "OR", ...)* function. Random and fixed effect models were fit using the rma function (method = "REML" and method = "FE", respectively). All necessary input data can be retrieved from supplementary table 6.

2. Simon Anders P, Wolfgang H. HTSeq—A Python framework to work with high-throughput sequencing data. Bioinformatics 2014.

3. Trapnell C, Williams BA, Pertea G, Mortazavi A, Kwan G, van Baren MJ, et al. Transcript assembly and quantification by RNA-Seq reveals unannotated

transcripts and isoform switching during cell differentiation. Nat Biotechnol 2010;28(5):511-5 doi 10.1038/nbt.1621.

4. Bailey P, Chang DK, Nones K, Johns AL, Patch A-M, Gingras M-C, et al. Genomic analyses identify molecular subtypes of pancreatic cancer. Nature 2016;531(7592):47-52 doi 10.1038/nature16965.

5. Wang L, Wang S, Li W. RSeQC: quality control of RNA-seq experiments. Bioinformatics 2012;28(16):2184-5 doi 10.1093/bioinformatics/bts356.

6. Love MI, Huber W, Anders S. Moderated estimation of fold change and dispersion for RNA-seq data with DESeq2. Genome Biol 2014;15(12):550 doi 10.1186/s13059-014-0550-8.

7. Team RC. R: A Language and Environment for Statistical Computing. 2016.

8. Fraley C, Raftery AE. MCLUST version 3: an R package for normal mixture modeling and model-based clustering. 2006.

9. Feldman J, Donnell RO, Servedio RA. Learning mixtures of product distributions over discrete domains. 2005 23-25 Oct. 2005. p 501-10.

10. Gaujoux R, Seoighe C. A flexible R package for nonnegative matrix factorization. BMC Bioinformatics 2010;11(1):367 doi 10.1186/1471-2105-11-367.

11. Schwender H. siggenes: Multiple testing using SAM and Efron's empirical Bayes approaches 2012.

24. Maechler M, Rousseeuw P, Struyf A, Hubert M, Hornik K. cluster: Cluster Analysis Basics and Extensions. 2017.

25. Therneau TM, Grambsch PM. Modeling Survival Data: Extending the Cox Model. Springer New York; 2013.

26. Freund Y, Schapire RE. A desicion-theoretic generalization of on-line learning and an application to boosting. InEuropean conference on computational learning theory 1995 Mar 13 (pp. 23-37). Springer, Berlin, Heidelberg.

27. Oh EY, Christensen SM, Ghanta S, Jeong JC, Bucur O, Glass B, Montaser-Kouhsari L, Knoblauch NW, Bertos N, Saleh SM, Haibe-Kains B. Extensive rewiring of epithelial-stromal co-expression networks in breast cancer. Genome biology. 2015 Jun 19;16(1):128.
