## Supplementary Materials for "Transcriptional deconvolution reveals consistent functional subtypes of pancreatic cancer epithelium and stroma"

### Supplementary Figure 1

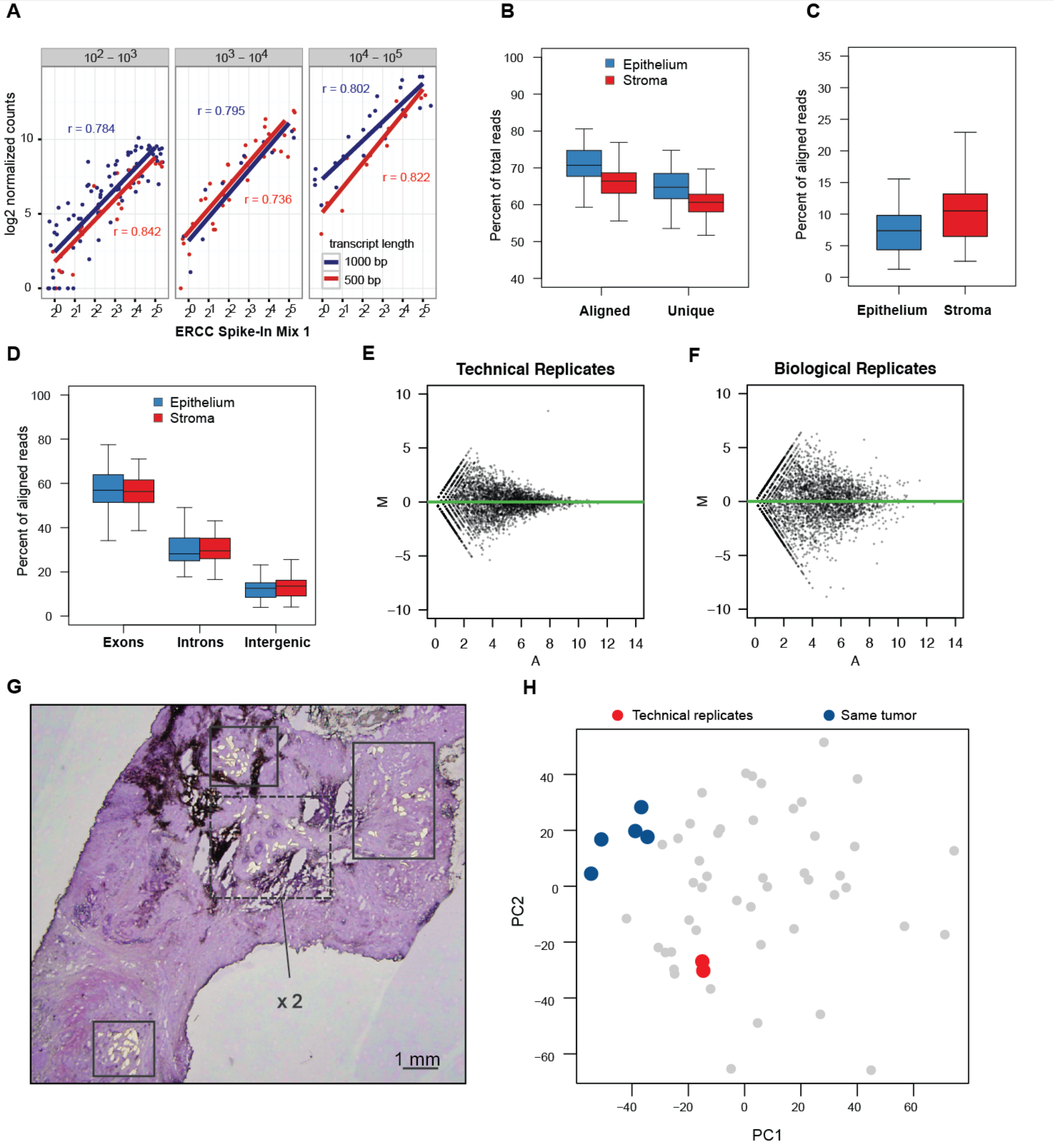


**Supplementary Figure 1. Quality control of the LCM-RNA-Seq pipeline**
(**A**) ERCC RNA Spike-In Mix 1 (Ambion) was added to LCM RNA samples before amplification at increasing concentrations. The amount of ERCC Spike-In Mix correlates strongly with the observed ERCC read counts across different concentration and length categories, suggesting that amplification of comparable mRNA species occurs linearly as well. After read alignment using Tophat (v.2.0.4), BAM files (n = 120, 60 per compartment) were analyzed using the RSeQC package (v.2.6.2) to quantify: (**B**) the fraction of aligned reads and uniquely aligned reads, respectively, as compared to the total number of reads (= 100%) generated for each library, (**C**) the fraction of reads mapping to ribosomal RNA genes, as compared to the number of aligned reads (= 100%). (**D**), the fraction of aligned reads aligning to exonic, intronic and intergenic regions, respectively. (**E**) MA-plots of technical vs. (**F**) biological replicates of laser captured epithelial tumor samples. (**G**) Cresyl violet stain of a frozen PDAC section on a PEN membrane slide. Boxes indicate areas from which epithelial cells were gathered using LCM. The area marked by the dashed line was sampled from both the section shown and a further serial section from this tumor with at least 50 µm distance. These 5 biological replicates (blue) from the same tumor cluster closely in (**H**) PCA when compared to further biological replicates from different tumors (grey) while technical replicates (red) show virtually identical behavior in PCA.
