## Supplementary Materials for "Transcriptional deconvolution reveals consistent functional subtypes of pancreatic cancer epithelium and stroma"

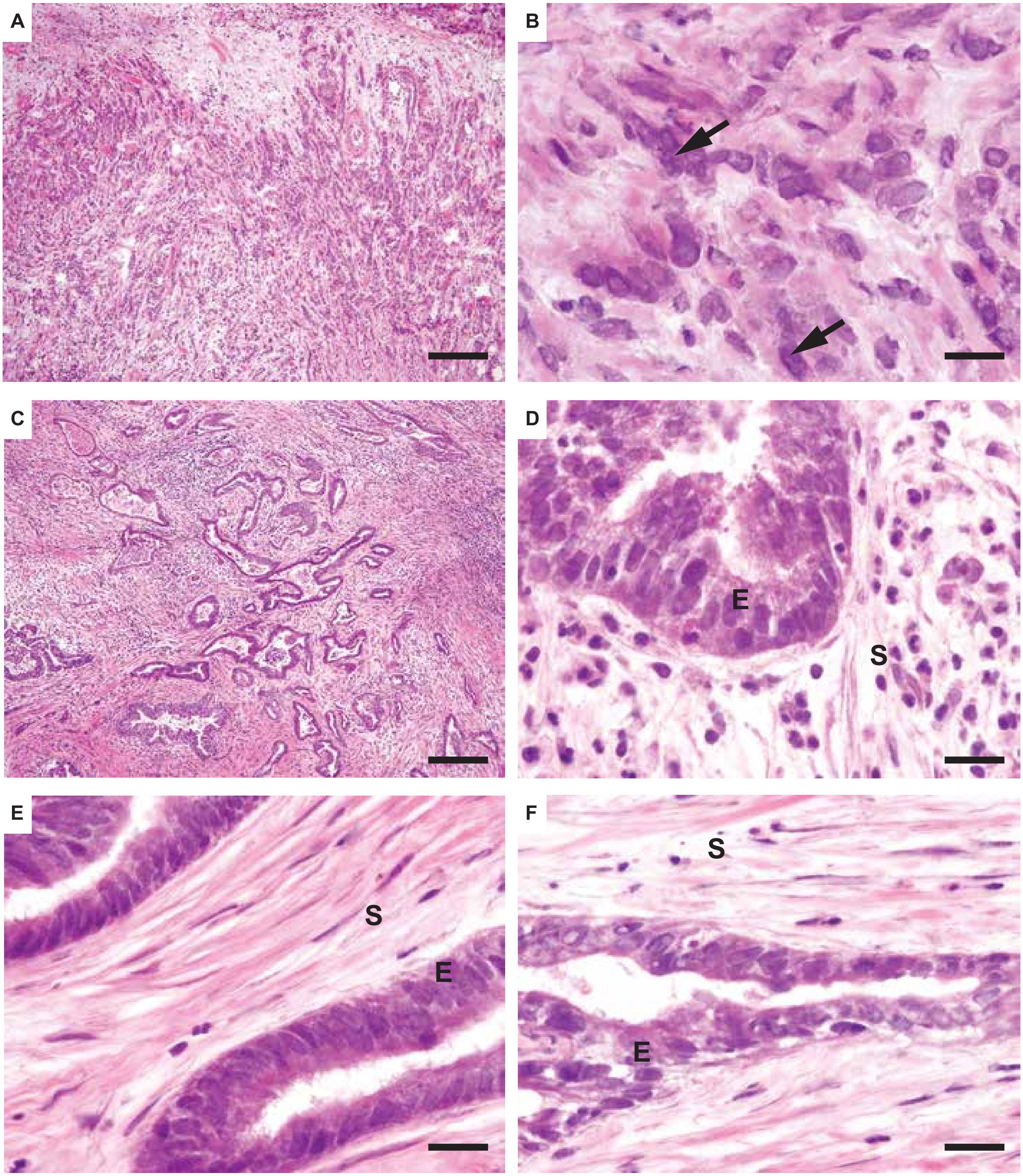


**Supplementary Figure 2. Histopathology of outlier samples.**
Images show hematoxylin and eosin stained sections from samples E17(A,B), S10 (C-E), and S7 (F). Scale bars for (A,C) indate 200μm and for (B,D,E,F) indicate 20μm. Epithelial cells in sample E17 (arrows) were poorly differentiated. Sample S10 exhibited stroma with two starkly different phenotypes: loosely packed, ECM-poor hypercellular stroma (D) and a more classical ECM-rich desmoplastic stroma (E). Sample S7 exhibited histopathology typical of PDA. E = epithelium; S = stroma.
