## Supplementary Materials for "Transcriptional deconvolution reveals consistent functional subtypes of pancreatic cancer epithelium and stroma"

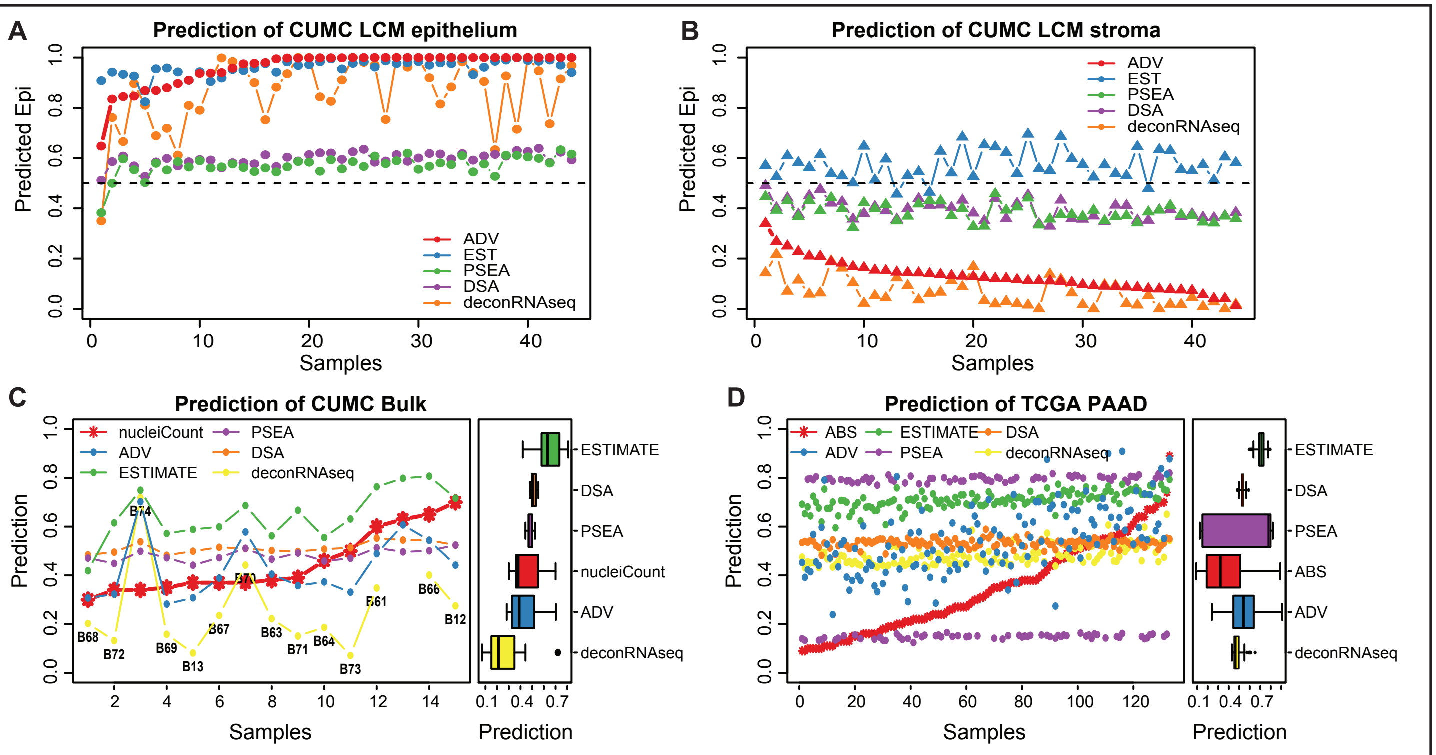


**E**

| | CUMC LCM samples | CUMC bulk samples | TCGA bulk samples |
|---|---|---|---|
| Dataset: | CUMC LCM samples | CUMC bulk samples | TCGA bulk samples |
| Reference data: | LCM epithelium | Nuclei counting | ABSOLUTE |
| | MAPE | MAPE | MAPE |
| ADVOCATE | 8.44 | 15.28 | 20.47 |
| ESTIMATE | 30.69 | 24.63 | 40.73 |
| PSEA | 40.76 | 15.63 | 38.02 |
| DSA | 39.89 | 17.04 | 24.93 |
| DeconRNASeq | 10.11 | 33.72 | 20.53 |

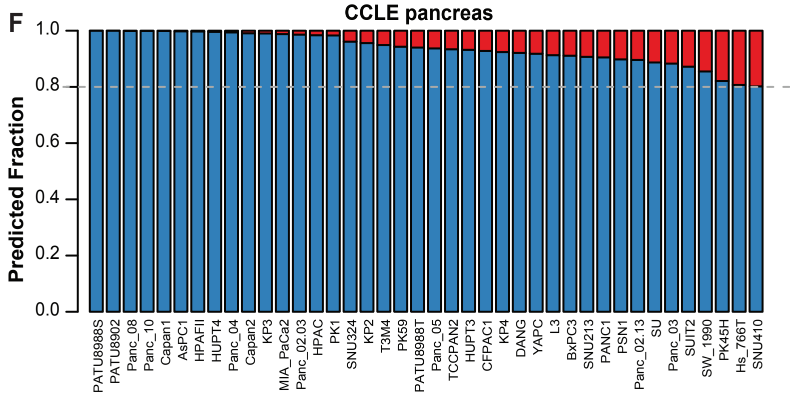


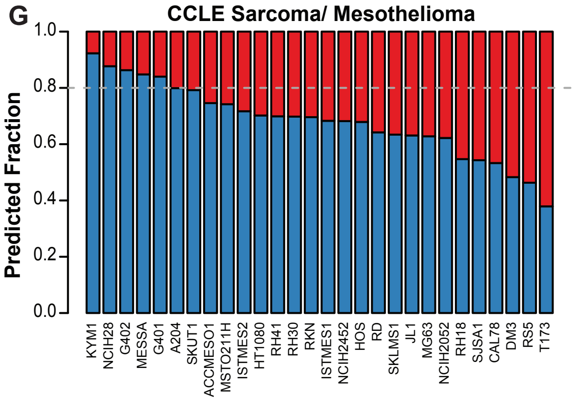


**Supplementary Figure 3. Algorithm and molecular extended**
(A-D) Comparison of tissue fraction predictions by different algorithms, using the (A) CUMC PDA LCM epithelial, (B) CUMC PDA LCM stromal, (C) CUMC bulk PDA, and (D) TCGA PDA bulk sample sets. Depicted as scatter plots by sample ID and boxplots (C)-(D) right. Algorithms are color-coded. X-axis for scatter plot is sample ID. Y-axis is algorithm predicted tumor purity. (E). Summary statistics of algorithm comparisons. MAPE: mean average percentage error. (F,G) ADVOCATE compartment fraction predictions for cell lines derived from (E) pancreatic ductal adenocarcinoma or (F) sarcomas and mesotheliomas. Expression profiles were obtained from the Cancer Cell Line Encyclopedia (CCLE).
