## Supplementary Materials for "Transcriptional deconvolution reveals consistent functional subtypes of pancreatic cancer epithelium and stroma"

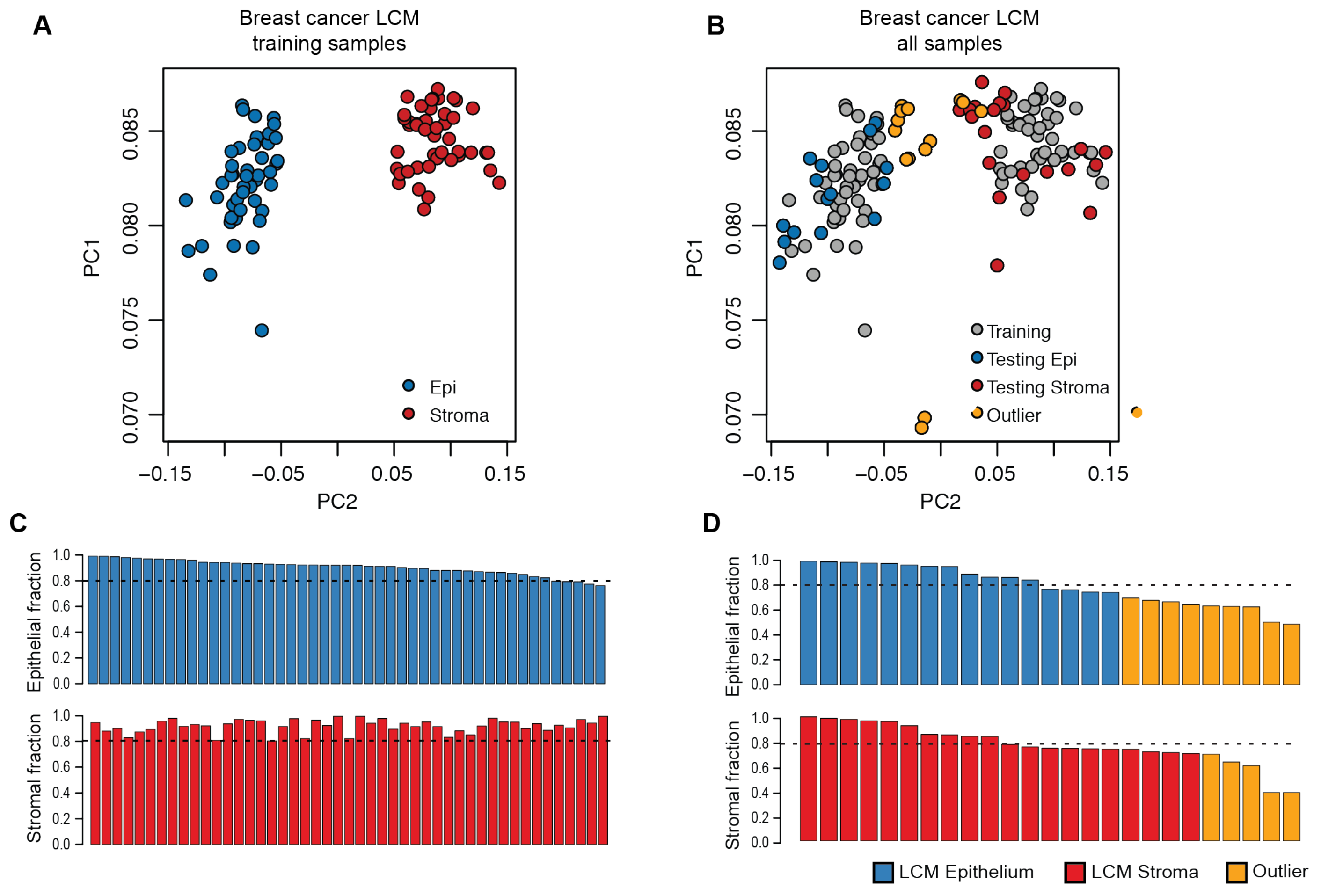


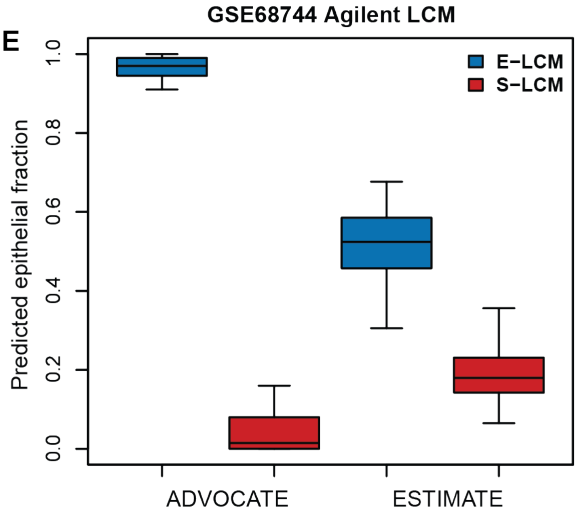

**Supplementary Figure 4. ADVOCATE implementation for breast cancer LCM data.** (**A**) PCA of 82 matched compartment-specific gene expression profiles from breast cancer samples as described by Oh et al. Compartment origin separates the samples along the second principal component. (**B**) PCA of the all LCM samples used for training a breast cancer implementation of ADVOCATE and independent validation LCM samples (colored points). (**C**) Results from a Leave-One-Out Cross-Validation (LOOCV). Epithelial samples were predicted to have high epithelial content (upper panel) and stromal samples were predicted to have high stromal content. (**D**) Predicted compartment fractions for an independent set of LCM samples (colored points in (B)), including outliers with little distinction between epithelium and stroma (yellow points in PCA plot). (**E**) Prediction of GSE68744 LCM Epithelium (E-LCM) and Stroma (S-LCM) Agilent profiles. Predictions from the implementation of ADVOCATE from (A) are compared to predictions from the ESTIMATE algorithm.
