## Supplementary Materials for "Transcriptional deconvolution reveals consistent functional subtypes of pancreatic cancer epithelium and stroma"

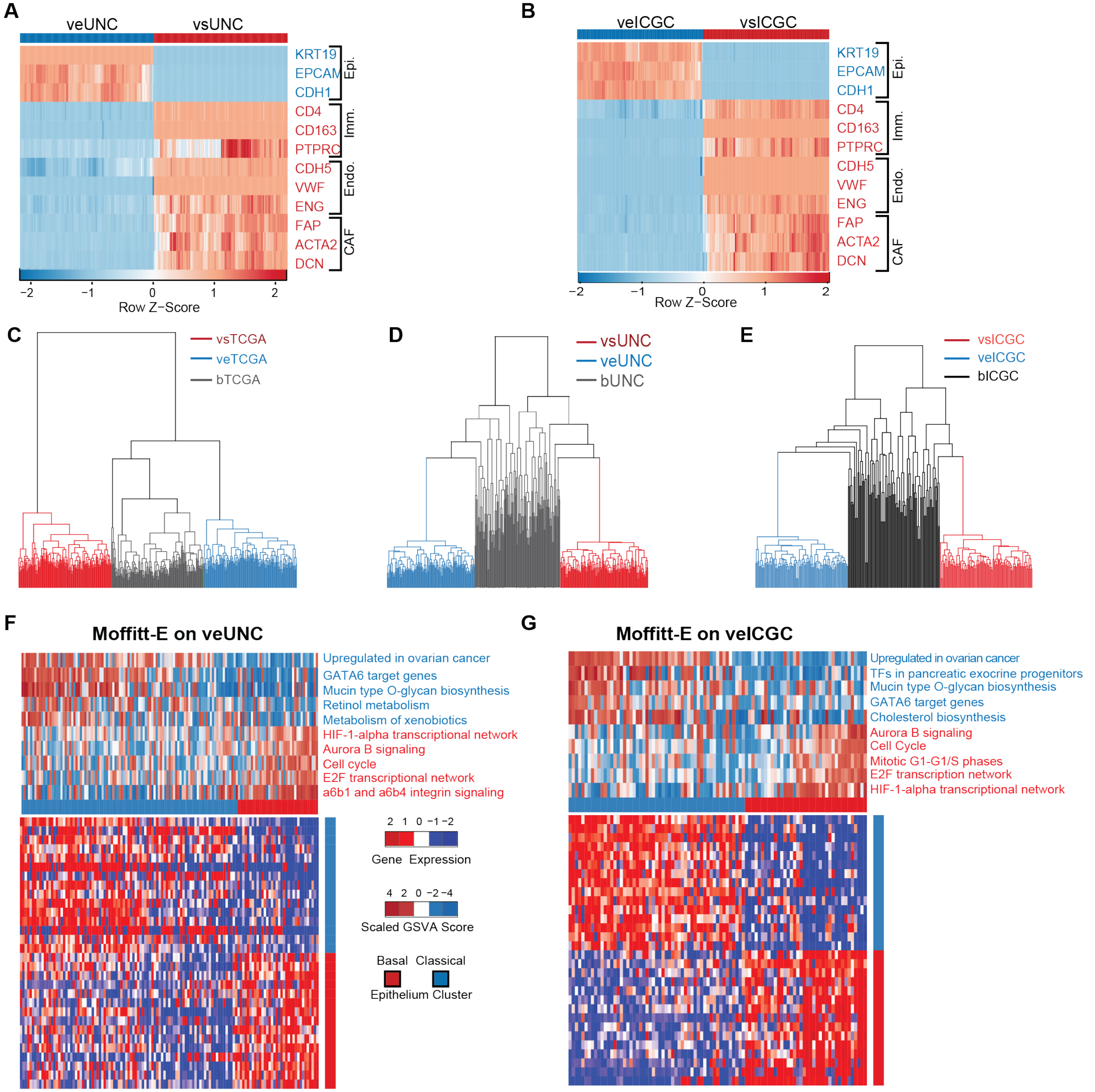


**Supplementary Figure 5. Deconvolution of external cohorts and epithelial subtyping**
(**A-B**) Heat-map showing the expression of indicated marker genes in deconvolved virtual epithelial and stromal profiles from the UNC (A) and ICGC (B) cohorts. Color scale indicates row Z-scores. (**C-E**) Hierarchical clustering of TCGA (C), UNC (D), and ICGC (E) bulk tumors (black) and their compartment–specific virtual derivatives (epithelial samples in blue, stromal samples in red). (**F-G**) Heatmaps of the top 30 differentially expressed genes between two groups that were obtained by clustering virtual epithelial UNC (F) and ICGC (G) profiles using the Moffitt–E classifier. GSVA scores for indicated gene set are represented at top. Tumors with Basal–like (red bar) and Classical traits (blue bar), respectively, can be detected in both virtual tumor cohorts.
