## Supplementary Materials for "Transcriptional deconvolution reveals consistent functional subtypes of pancreatic cancer epithelium and stroma"

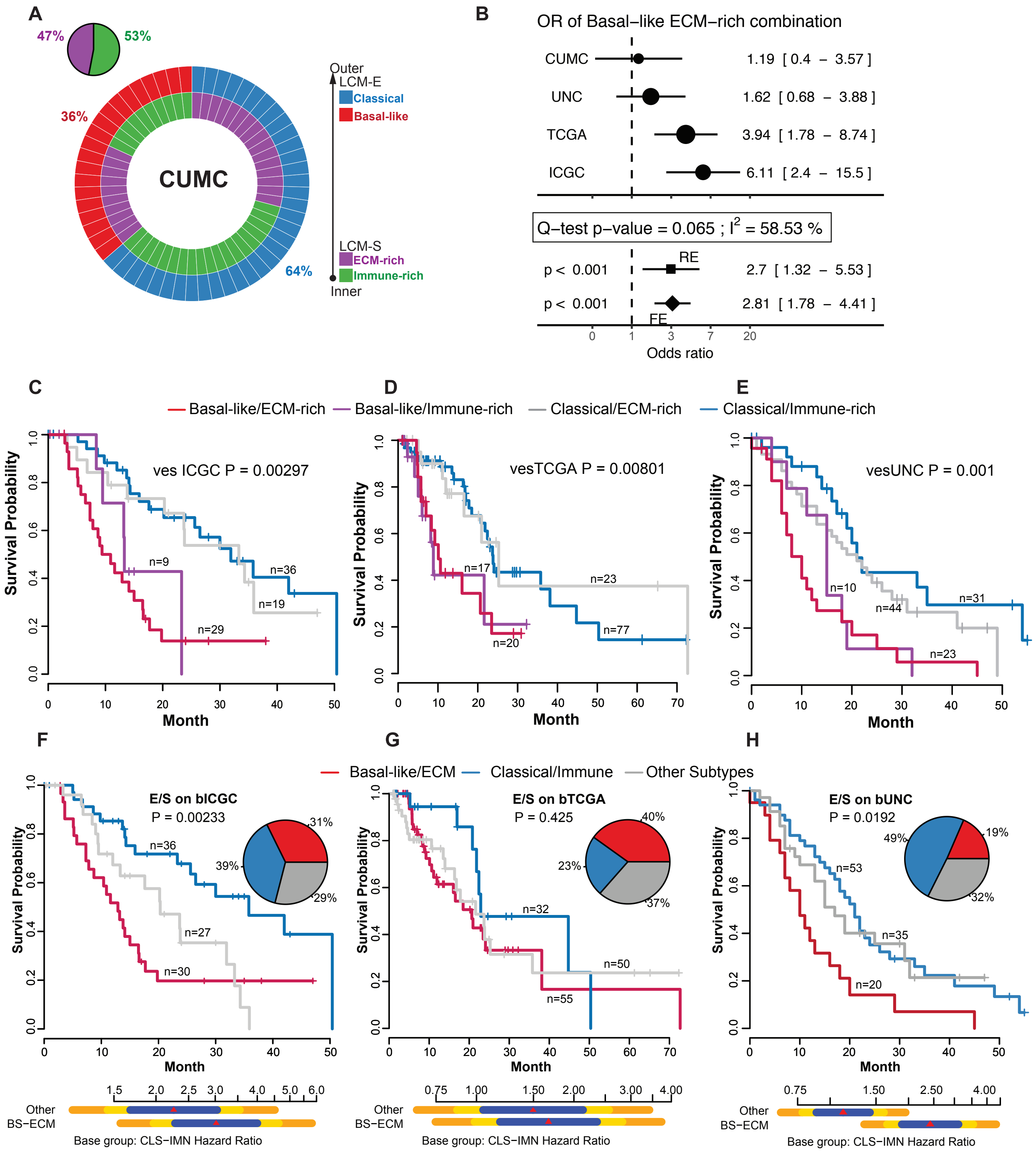


**Supplementary Figure 6. Bi-Compartmental subtyping of PDA extended**
(**A**) Multilayered donut plot showing: (i) the alignment of epithelial and stromal subtypes for LCM profiles from CUMC tumors, and (ii) the proportion of each epithelial subtype. Separate pie chart summarizes the proportion of stromal subtypes. (**B**) Odds ratios of Basal-like tumors having ECM-rich stromas across the indicated cohorts including their 95% confidence interval (upper panel). Meta-analytic fixed (FE)- and random-effects (RE) models show significant positive associations. Q-test and $I^2$-statistic indicate measures of inter-study heterogeneity (**C-E**) KM survival analysis per combined subtype as classified from deconvolved GEP from the indicated cohorts. P-values reflect the probability that all groups have the same survival function. (**F-H**) KM survival analysis of combined epithelial and stromal subtypes as determined from bulk expression profiles for each of the indicated cohorts. Red and blue lines contrast Basal–like/ECM–rich tumors against Classical/Immune–rich tumors while all other tumors are represented as a grey line. Pie charts summarize the proportion of each category per cohort. In contrast to the combined subtyping results that were obtained using virtual compartment–specific profiles, the Basal–like/ECM–rich subtype is not associated with a significantly reduced survival in the bulk TCGA cohort.
